# Estimating the Contribution of Proteasomal Spliced Peptides to the HLA-I Ligandome

**DOI:** 10.1101/288209

**Authors:** Roman Mylonas, Ilan Beer, Christian Iseli, Chloe Chong, HuiSong Pak, David Gfeller, George Coukos, loannis Xenarios, Markus Müller, Michal Bassani-Sternberg

**Affiliations:** Vital-IT, Swiss Institute of Bioinformatics, 1015 Lausanne, Switzerland.; Swiss Institute of Bioinformatics, 1015 Lausanne, Switzerland.; Adicet Bio Israel Ltd, Technion City, 32000, Haifa, Israel.; Ludwig Cancer Research Center, University of Lausanne, 1066 Epalinges, Switzerland.; Department of Oncology, University Hospital of Lausanne, 1011 Lausanne, Switzerland.

**Keywords:** de-novo sequencing, HLA, Immunopeptidomics, proteasomal splicing

## Abstract

Spliced peptides are short protein fragments spliced together in the proteasome by peptide bond formation. True estimation of the contribution of proteasome-spliced peptides (PSPs) to the global Human Leukocyte Antigen (HLA) ligandome is critical. A recent study suggested that PSPs contribute up to 30% of the HLA ligandome. We performed a thorough reanalysis of the reported results using multiple computational tools and various validation steps and concluded that only a fraction of the proposed PSPs passes the quality filters. To better estimate the actual number of PSPs, we present an alternative workflow. We performed de-novo sequencing of the HLA-peptide spectra and discarded all de-novo sequences found in the UniProt database. We checked whether the remaining de-novo sequences could match spliced peptides from human proteins. The spliced sequences were appended to the UniProt fasta file, which was searched by two search tools at a FDR of 1%. We find that maximally 2-4% of the HLA ligandome could be explained as spliced protein fragments. The majority of these potential PSPs have good peptide-spectrum match properties and are predicted to bind the respective HLA molecules. However, it remains to be shown how many of these potential PSPs actually originate from proteasomal splicing events.

**Abbreviations**
APPM: Antigen processing and presentation machinery
HLA: Human Leukocyte Antigen
HLA-Ip: HLA class I binding peptides
FDR: False discovery rate; ***FP***: false positive
AA: amino acid
LC: liquid chromatography
MS: Mass spectrometry
MS/MS: Tandem mass spectrometry
PSPs: *Proteasome-spliced peptides*
DeNovo_spliced: Spliced peptides identified by de-novo
*DeNovo_non-spliced*: Non-spliced peptides not in UniProt identified by de-novo
LM_spliced: Spliced peptides identified by Liepe et al.
LM_UniProt: UniProt peptides identified by Liepe et al.
PSM: Peptide spectrum match

## Introduction

The antigen processing and presentation machinery (APPM) is responsible for the cell surface display of thousands of peptides in the context of the HLA class I (HLA-I) molecules. The proteasome is considered as the main protease that cleaves endogenous proteins. However, in addition to the proteasome, the APPM comprises several other proteases, transporters and chaperones that cooperatively digest the proteins in the cytoplasm, funnel the peptides into the ER, further trim and edit them, load them on newly synthesized HLA-I, and finally direct the stable complexes to the cells’ surface (1). The selective interaction between the HLA-I complex and the peptides is the major factor that defines the presented repertoire and is often represented with binding motifs.

Currently, the only unbiased methodology to comprehensively interrogate the repertoire of the HLA-I binding peptides (HLA-Ip) is based on mass spectrometry (MS). HLA complexes are immunoaffinity-purified from cells in culture or from tissues; the peptides are extracted and subjected to reverse-phase liquid chromatography (LC) coupled online to sensitive MS instruments. The acquired tandem mass spectrometry (MS/MS) data is normally searched against a database of protein sequences. Applying a stringent FDR of 1% using a comparable decoy database leads to the accurate identification of thousands of HLA-Ip. HLA-Ip are mainly 9-11 amino acids (AA) long and usually about 95% of the peptides identified with this methodology fit the consensus binding motifs of the HLA expressed in the samples (2).

In a recent MS-based HLA-I ligandomics study a novel computational algorithm has predicted that a surprisingly large fraction, up to 30%, of the ligands may be derived from transpeptidation of two noncontiguous fragments of a parental protein that are spliced together within the proteasome (3). Earlier work showed several cases of such proteasomal spliced HLA-I peptides that were naturally presented and recognized by cytotoxic T cells (4-9). Hence, these may be highly interesting therapeutic targets. However, the authors of (3) noticed that unlike the non-spliced peptides, proteasome-spliced peptides (PSPs) had low HLA binding affinities and produced ambiguous binding motifs compared to normal HLA-Ip. HLA loading takes place after the peptides have exited the proteasome and entered the ER, and hence lost the identity of their creation mechanism. Currently, there is no mechanism or biological process that could explain how the APPM can prioritize loading of HLA-I molecules with low affinity PSPs over high-affinity non-spliced peptides.

Understanding the contribution of PSPs to the HLA ligandome is crucial, especially as they may indeed be highly interesting therapeutic targets in many diseases. Here we critically investigated PSPs reported in Liepe et al. (3) and found that most of spectra attributed to them could be assigned with higher scores to normal peptide sequences within the UniProt database of human proteins. We further describe an alternative computational pipeline to estimate the contribution of spliced peptides to the immunopeptidome. Our results suggest that less than 2-4% of the HLA-Ip may be spliced. As opposed to the spliced peptides reported in (3), these peptides fit well to the relevant HLA binding motifs.

## Experimental Procedures

### HLA ligandomic data

We selected previously published MS HLA-Ip datasets of exceptionally high coverage representing a variety of binding specificities (Supplemental Table 1). MS raw files of HLA-Ip isolated from two melanoma tissues, Mel15 (16 raw files) and Mel16 (12 raw files) (10), RA957 B cell line (4 raw files) (11) and Fibroblast (Fib) cells (4 raw files) (2) were downloaded from the PRIDE repository (12) dataset PXD004894, PXD005231 and PXD000394, respectively. One of the four raw MS files of the Fib cells (20130504_EXQ3_MiBa_SA_Fib-2.raw) was also used by Liepe et al. More details about these datasets can be found on PRIDE and the respective manuscripts.

### Data processing

If not otherwise mentioned, data were processed with the R statistical scripting language (version 3.3.2) (https://www.r-project.org/).

## Experimental design and statistical rationale

### Identification of HLA-Ip using PEAKS

Raw files were analyzed with the de-novo sequencing software PEAKS Studio 8.0 (13). General parameters were set to “Ion Source”: ESI (nano-spray), “Fragmentation Mode”: high energy CID (y and b ions), “MS Scan Mode” and “MS/MS Scan Mode”: FT-ICR/Orbitrap. The different PEAKS modules were used in the following order with their default parameters while special parameters are indicated in parenthesis: 1. DATA REFINE, 2. DENOVO (“Parent Mass Error Tolerance”: 10 ppm, “Fragment Mass Error Tolerance”: 0.02 Da, “Enzyme”: None), 3. PEAKS (“Parent Mass Error Tolerance”: 10 ppm, “Fragment Mass Error Tolerance”: 0.02 Da, “Variable Modifications”: Oxidation (M) 15.99, “Database”: Homo_sapiens_UP000005640_9606), 4. PEAKS PTM (“Parent Mass Error Tolerance”: 10 ppm, “Fragment Mass Error Tolerance”: 0.02 Da, “Enzyme”: None, “Database”: Homo_sapiens_UP000005640_9606), 5. SPIDER (“Variable Modifications”: Oxidation (M) 15.99, “Fragment ion tolerance”: 0.02). All peptides with -10LogP > 15 were considered having a match with a human protein from UniProt (Homo_sapiens_UP000005640_9606). Peptides with a -10LogP ≤15 (FDR around 1.5%) and a ALC% ≥ 15 were considered as “de novo only peptides”. Peptides with mutations and modifications were ignored.

For the subsequent analysis we merely kept “de novo only peptides” with a length between 8 and 25 AA. For each PSM we kept the highest ranked match with a local confidence score over 80 for every amino acid position if not otherwise mentioned. In order to simplify association of peptides with their corresponding HLA alleles, peptides containing PTMs were removed.

### Identification of possible spliced-peptides using TagPep

We checked whether the filtered list of sequences from the “de-novo only peptides”, which did not match any UniProt sequence, could be spliced fragments from UniProt (Homo_sapiens_UP000005640_9606) proteins. We applied an in-house software tool TagPep, which uses the index strategy described for fetchGWI (14) adapted for AAs instead of nucleotides. TagPep first matches the whole peptide sequence to the database. If there is no complete hit, it looks for hits allowing for one splicing event, where both spliced fragments are from the same protein (Supplemental Data 1). We excluded *trans*-spliced peptides where the fragments stem from two different proteins for three reasons. First, all spliced peptides reported in (15) are concatenated fragments from the same protein. Second, the huge number of *trans*-spliced may lead to strongly increased false positive rates in subsequent bioinformatics analysis, and third, for trans-splicing to happen the two source proteins need to be present in the same proteasome at the same time, which is unlikely to happen on a large scale. The spliced fragments can lie anywhere in the protein, but their sequences cannot overlap. We only considered TagPep matches if the splice gap, i.e. the minimal distance between the two spliced fragments, was less than 20 AA. It has been argued that the splice gap in truly existing spliced peptides is usually lower than 20 AAs (16). This distance threshold was used in Liepe et al. (3). We named the resulting set of spliced peptides as *DeNovo_spliced*. The remaining peptides in the PEAKS “de-novo only peptides” list that are not present in *DeNovo_spliced* were named *DeNovo_non-spliced*.

Of note, PEAKS de-novo assigns the mass 113.08406 by default as Leucine, and therefore the TagPep results may bias against identification of Isoleucine containing sequences. These isobaric amino acids make up 10% of all amino acids. However, even if 50% of the Leucine/Isoleucine PEAKS assignments are misplaced, this would lead to a wrong TagPep match in roughly 5-10% of all spliced peptides (assuming one or two Leucine/Isoleucine per peptide); therefore, will not significantly change our main results.

### Confirmation of identification of spliced peptides using MaxQuant and Comet

We employed the MaxQuant platform (17) version 1.5.5.1 and the Comet software release 2016.01 (18) to search the peak lists against the UniProt database (Homo_sapiens_UP000005640_9606) and a fasta file containing 247 frequently observed contaminants. For each sample, we added to the fasta file the list of *DeNovo_spliced* and the *DeNovo_non-spliced* peptide sequences. For Fib, we also added the 1,154 spliced peptides identified by Liepe et al. (3), which we named *LM_spliced*. The list of spliced peptides was kindly provided to us by the authors of (3) (Supplemental Data 2, type = “psp”). Peptides with a length between 8 and 25 AA were allowed. MaxQuant parameters: The second peptide identification option in Andromeda was enabled. The enzyme specificity was set as unspecific. A FDR of 1% was required for peptides and no protein FDR was set. The initial allowed mass deviation of the precursor ion was set to 6 ppm and the maximum fragment mass deviation was set to 20 ppm. Only peptides identified with delta score larger than zero were considered. Methionine oxidation (15.994915 Da) and N-terminal acetylation (42.010565 Da) were set as variable modifications, however modified peptides identified in Fib cells were removed at first for the direct comparison to the Liepe's data. For the additional modification search, methionine oxidation N-terminal acetylation, as well as glutamine/asparagine deamidation (+0.98402 Da) were set as variable modifications. Comet parameters: activation method: HCD, peptide mass tolerance: 0.02 Da, fragment mass tolerance: 0.02, fragments: b- and y-ions, precursor tolerance, modifications: for the modification search same as MaxQuant plus methylation (14.01565 Da) on K,R,D,E.

### LC-MS/MS analyses and identification of selected synthetic HLA-Ip

Synthetic peptides (PEPotech Heavy grade 3, Thermo Fisher Scientific) (listed in Supplemental Table 2) corresponding to peptides identified from Fib data were mixed and desalted on a C-18 spin column (Harvard Apparatus, 74-4101) and measured at a total amount of 10 and 20 pmol. Synthetic peptides were separated by a nanoflow HPLC (Proxeon Biosystems, Thermo Fisher Scientific, Odense) and coupled on-line to a Q Exactive HF mass spectrometer (Thermo Fisher Scientific, Bremen) with a nanoelectrospray ion source (Proxeon Biosystems). We packed a 20 cm long, 75 μm inner diameter column with ReproSil-Pur C18-AQ 1.9 μm resin (Dr. Maisch GmbH, Ammerbuch-Entringen, Germany) in buffer A (0.5% acetic acid). Peptides were eluted with a linear gradient of 2-30% buffer B (80% ACN and 0.5% acetic acid) at a flow rate of 250 nl/min over 90 min. Data was acquired using a data-dependent ‘top 10’ method. We acquired full scan MS spectra at a resolution of 70,000 at 200 m/z with an Auto gain control (AGC) target value of 3e6 ions. Ten most abundant ions were sequentially isolated, activated by Higher-energy collisional dissociation and accumulated to an AGC target value of 1e5 with a maximum injection time of 120 ms. In case of unassigned precursor ion charge states, or charge states of four and above, no fragmentation was performed. The peptide match option was disabled. MS/MS resolution was set to 17,500 at 200 m/z. Selected ions form fragmentation were dynamically excluded from further selection for 20 seconds. We employed the MaxQuant settings mentioned above for synthetic peptides identification.

### Comparison of MS/MS annotations of endogenous HLA-Ip and their synthetic counterparts

To investigate if spliced peptides match the MS/MS spectra better than possible alternative sequences we first compared the MS/MS scans identified by Liepe et al. as spliced peptides and those identified by MaxQuant. The mapping to the relevant MS scans and their Mascot ion scores were kindly provided to us by the authors of (3) (Supplemental Data 2). For the Fib data, we selected three MS/MS scans of *LM_spliced* peptides identified both by MaxQuant and by Liepe et al., and 21 MS/MS scans corresponding to *LM_spliced* peptides that MaxQuant identified instead as UniProt peptides. This selection was not biased and was not based on prior knowledge, which would favor MaxQuant. We synthesized the 21 pairs of peptide sequences and the three *LM_spliced* peptide sequences and analyzed them by MS as mentioned above. For visual inspection we printed the endogenous and synthetic spectra to pdf files. For each of the 21 pairs, we calculated the similarity between the spectrum of the eluted peptide from Fib, annotated as *LM_spliced* and the spectrum of the synthetic *LM_spliced* peptide. We also calculated the similarity between the spectrum of the eluted peptide from Fib annotated as a UniProt peptide, and the spectrum of the synthetic Uniprot peptide. Similarly, we calculated the similarity between the three spectra of the identically identified spliced sequences and the spectra of their synthetic counterparts. The similarity was computed by the cosine score (correlation coefficient; value between 0 and 1, where a value of 0 corresponds to spectra with no peaks in common and a value of 1 to identical spectra) (19). The MzJava class library (20) was used to read the .mgf spectrum files and to calculate the similarity.

### Binding affinity prediction and clustering of peptides

We used the NetMHCpan (21) to predict binding affinity of 8-14 mer peptides to the respective HLA alleles expressed in the sample and assigned them based on maximum affinity. Hits with a rank <2% were considered as binders. Gibbscluster-1.1 (22) was run independently for each list of peptides identified from the different samples, with the default settings except that the number of clusters was tested between 1 and 6, a trash cluster was enabled and alignment was disabled (23). The MixMHCp tool (http://mixmhcp.org/) was used to cluster the peptides with default settings (11, 24).

## Results

### Predicted spliced peptides from Liepe et al. do not fit well to the consensus binding motifs

Spliced HLA-Ip identified by Liepe et al. in the Fib sample *(LM_spliced)* were reported to be barely compatible with the corresponding HLA-I binding motifs (3). First, we tested to what extent the reported 1,154 *LM_spliced* peptides *LM_spliced* follow the same binding specificity as the other 2,882 identified HLA-Ip, which matched proteins in UniProt *(LM_UniProt*, type = “pcp” in Supplemental Data 2). 90% of the *LM_UniProt* peptides were predicted binders by NetMHCpan compared to only 33% of the *LM_spliced* peptides (Supplemental Figure 1 A and B). Second, we used the computational tools MixMHCp (11, 24) and GibbsCluster (22) to identify the consensus binding motifs within the lists of 9-mer HLA-Ip in a fully unsupervised way (i.e., without predicting their binding affinity). Four dominant motifs corresponding to the HLA-A and HLA-B allotypes expressed in the Fib cells were identified for the *LM_UniProt* peptides with both methods, whereas the motifs found in the *LM_spliced* peptides were much less specific and did not match the known alleles (Figure 1 A). Furthermore, we observed a different length distribution of the *LM_spliced* and *LM_UniProt* peptides (Figure 1 B). *LM_Uniprot* peptides followed the expected peptide length distributions of HLA-I alleles with the majority of peptides of length 9, while *LM_spliced* peptides unexpectedly displayed the same amount of 9- and 10-mers. We obtain similar results for the fraction of 1,583 spliced and 3,779 non-spliced 9 mer peptides of the GR-LCL 2D data reported by Liepe et al. (Supplemental Figure 1 C and D). Altogether, our results show that expected HLA-Ip characteristics can be clearly recovered from the list of UniProt peptides, but cannot be observed in the spliced peptides.

**Figure 1.**
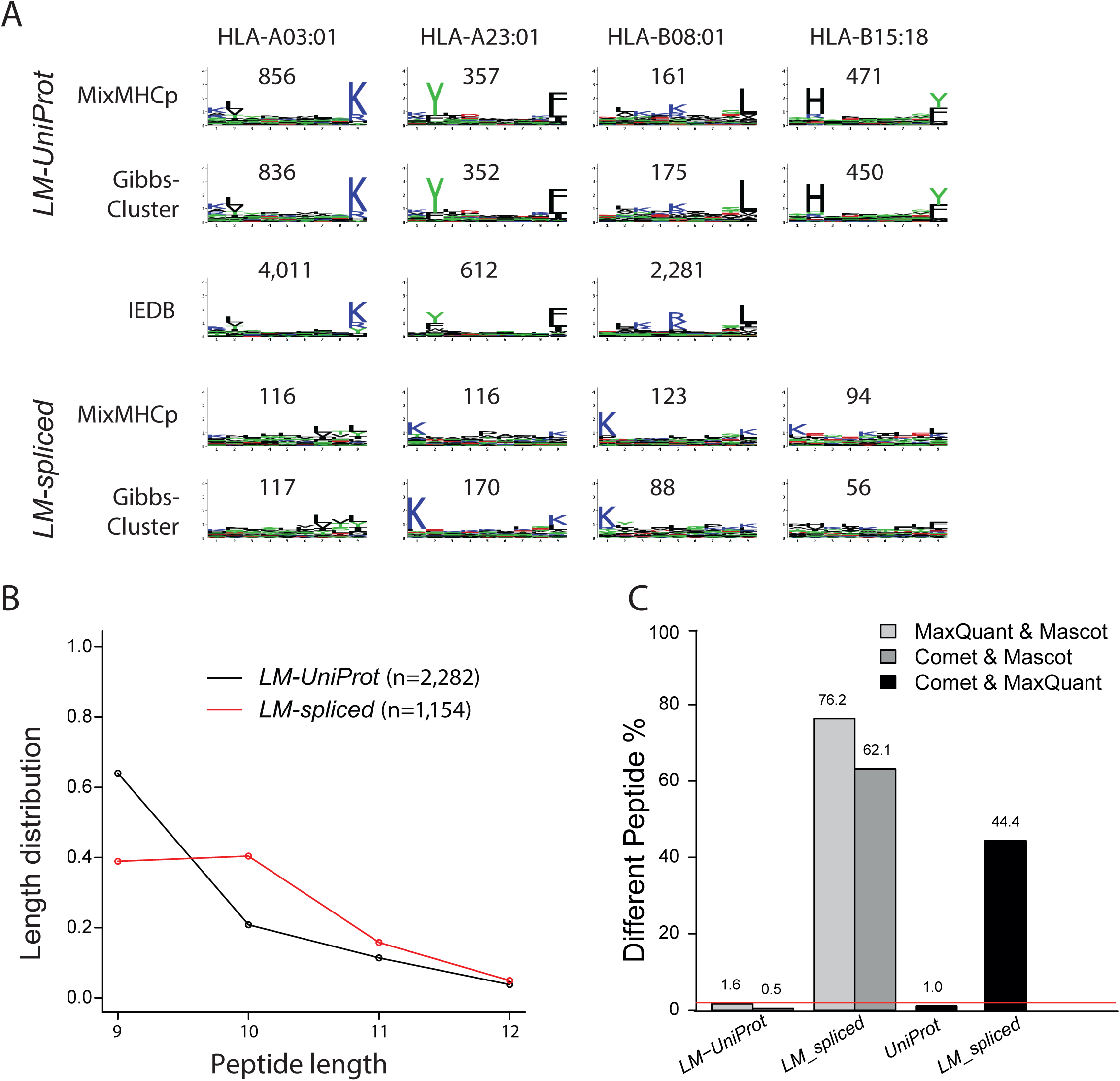
Motif deconvolution analysis with MixMHCp and GibbsCluster of the 9-mer *LM-UniProt* and *LM-spliced* peptides, and comparison to known logos from IEDB for HLA-A03:01, HLA-A23:01 and HLA-B08:01 (HLA-B15:18 has no experimental ligands in IEDB). Motifs found in *LM-Uniprot* peptides are highly reproducible and comparable to the known motifs from IEDB, while this is not the case for motifs found in *LM-spliced* peptides. B) Length distribution of the *LM-UniProt* and *LM-spliced* peptide. C) Rate of differing PSMs for the peptides in *LM-UniProt* and *LM-spliced*. Only MS/MS scans where both search strategies reported a match at FDR of 1% were considered.

### Spliced peptides from Liepe et al. produce more ambiguous peptide spectrum matches (PSMs)

We checked whether the spliced PSMs reported by Liepe et al. could be explained as matches to UniProt peptides. We added the list of 1,154 *LM_spliced* peptide sequences to the UniProt fasta file and researched the MS/MS data of the Fib raw file used by Liepe et al. using MaxQuant and Comet firstly without considering variable modifications. Out of the 6,358 (7,523) MS/MS scans identified by MaxQuant (Comet) at an FDR of 1%, 3,318 (3,522) MS/MS scans were also matched formerly by Liepe et al. (Table 1). Regarding these common MS/MS, we found very good agreement between MaxQuant and Comet in terms of matches of MS/MS scans to *LM_UniProt*. Only 1.6% (0.5%) of these scans matched a different peptide sequence in the MaxQuant (Comet) search (Figure 1 C). Since all searches were performed at a spectrum level FDR of 1%, the differences should not be larger than 2% and these values are within this range. However, for the group of common MS/MS scans, which matched to *LM_spliced* peptides in (3), the disagreement was 76.2% and 62.1% for MaxQuant and Comet, respectively. MaxQuant or Comet matched a UniProt peptide for the majority of these conflicts (98.2% for MaxQuant and 98.3% for Comet) and only in a few cases a different spliced peptide sequences. Overall, MaxQuant could confirm identification of 144 out of the 1,154 (12.4%) spliced peptides found by Liepe et al., whereas Comet could confirm 179 (15.5%). Therefore, the results obtained by Liepe et al., where the spliced peptides scored higher than the competing UniProt peptides, could not be confirmed by this reanalysis. More details about these comparisons can be found in Table 1. Interestingly, if we compare results between MaxQuant and Comet, the group of MS/MS scans matched to a peptide in *LM_spliced* also show a highly inflated disagreement (44.4%) compared to MS/MS scans matched to UniProt (1.0%) (Figure 1 C). Therefore, these results do not depend on the particular choice of search tool, but it seems that the PSMs assigned as *LM_spliced* peptides were more ambiguous compared to UniProt peptides, possibly because many *LM_spliced* peptides bear strong similarity to UniProt peptides.

**Table 1:**
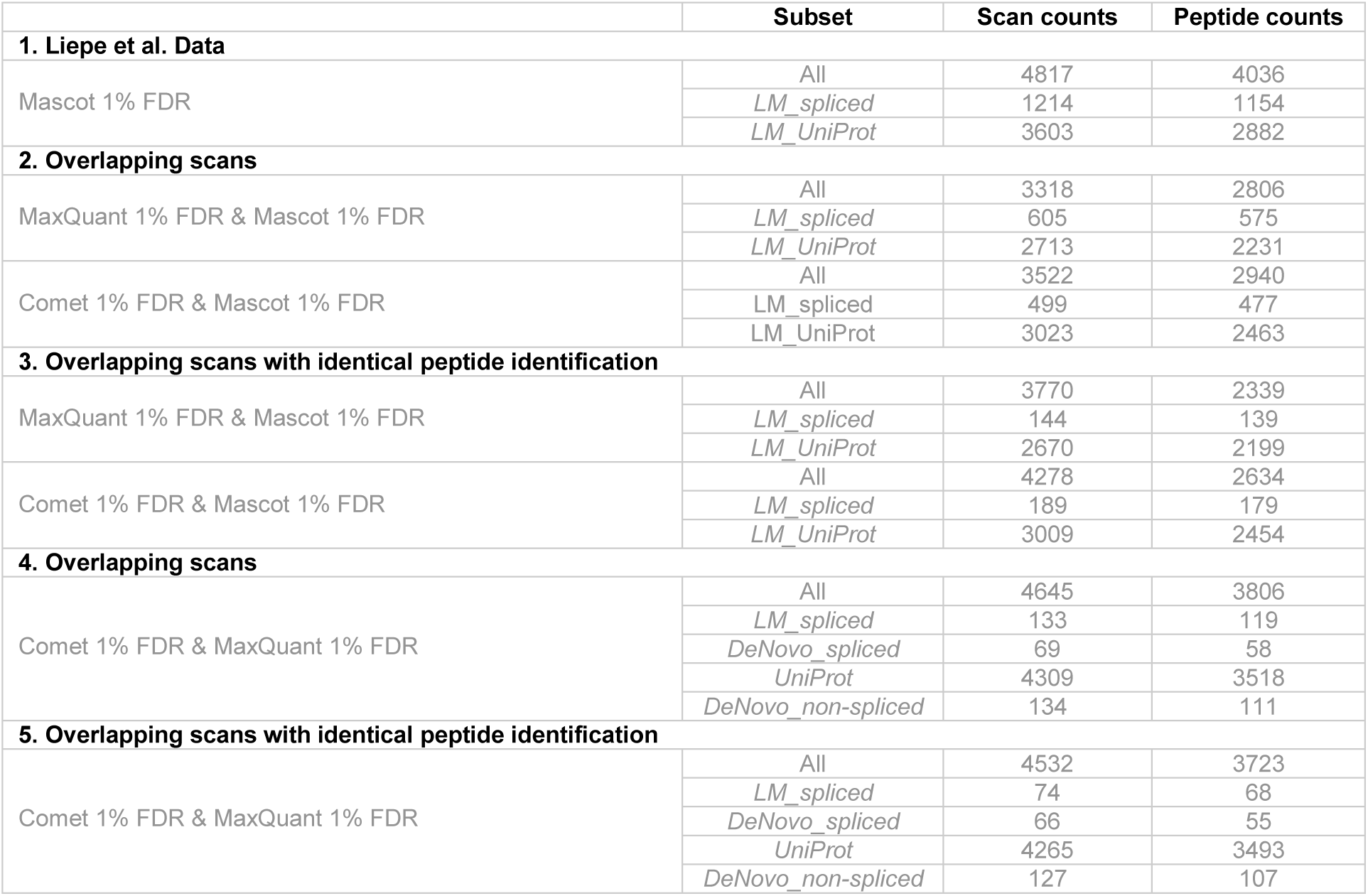
Summary of the level of agreement in scan matching and peptide identifications between Mascot (from Liepe et al.), MaxQuant and Comet for the different subset of peptides.

|  | Subset | Scan counts | Peptide counts |
| --- | --- | --- | --- |
| **1. Liepe et al. Data** |  |  |  |
| Mascot 1% FDR | All | 4817 | 4036 |
|  | *LM_spliced* | 1214 | 1154 |
|  | *LM_UniProt* | 3603 | 2882 |
| **2. Overlapping scans** |  |  |  |
| MaxQuant 1% FDR & Mascot 1% FDR | All | 3318 | 2806 |
|  | *LM_spliced* | 605 | 575 |
|  | *LM_UniProt* | 2713 | 2231 |
| Comet 1% FDR & Mascot 1% FDR | All | 3522 | 2940 |
|  | *LM_spliced* | 499 | 477 |
|  | *LM_UniProt* | 3023 | 2463 |
| **3. Overlapping scans with identical peptide identification** |  |  |  |
| MaxQuant 1% FDR & Mascot 1% FDR | All | 3770 | 2339 |
|  | *LM_spliced* | 144 | 139 |
|  | *LM_UniProt* | 2670 | 2199 |
| Comet 1% FDR & Mascot 1% FDR | All | 4278 | 2634 |
|  | *LM_spliced* | 189 | 179 |
|  | *LM_UniProt* | 3009 | 2454 |
| **4. Overlapping scans** |  |  |  |
| Comet 1% FDR & MaxQuant 1% FDR | All | 4645 | 3806 |
|  | *LM_spliced* | 133 | 119 |
|  | *DeNovo_spliced* | 69 | 58 |
|  | *UniProt* | 4309 | 3518 |
|  | *DeNovo_non-spliced* | 134 | 111 |
| **5. Overlapping scans with identical peptide identification** |  |  |  |
| Comet 1% FDR & MaxQuant 1% FDR | All | 4532 | 3723 |
|  | *LM_spliced* | 74 | 68 |
|  | *DeNovo_spliced* | 66 | 55 |
|  | *UniProt* | 4265 | 3493 |
|  | *DeNovo_non-spliced* | 127 | 107 |

To increase the coverage of possible spliced peptides, we searched three additional raw files of HLA-Ip derived from the same Fib cells. In total, MaxQuant identified 202 (17.5 %) and Comet 213 (18.5%) peptides as *LM_spliced* sequences (Supplemental Table 3). Compared to PSMs assigned as UniProt, peptides assigned as *LM_spliced* were characterized by lower Andromeda score for the best MS/MS spectrum (Supplemental Figure 2 A), lower Andromeda score difference to the second best identified peptide (Supplemental Figure 2 B), higher absolute precursor mass deviation (Supplemental Figure 2 C), fewer peaks matching to the predicted fragmentation spectrum (Supplemental Figure 2 D), lower fraction of total MS/MS peak intensity matched (Supplemental Figure 2 E) and a larger fraction of singly charged MS/MS spectra matched (Supplemental Figure 2 F). In agreement to our previous analysis employing a single Fib raw file, the combined analysis of four files showed similar results; the group of MS/MS scans matched to a spliced peptide from Liepe et al. showed a highly inflated disagreement (43.1%) compared to the UniProt scans (0.9%) (Supplemental Figure 2 G). Altogether, these results indicate the matches to most *LM_spliced* peptides are of lower quality.

### Spliced peptides from Liepe et al. conflict with modified UniProt peptides

Next, we tested how the inclusion of variable modifications in the searches influences the identification rates in the different groups. When adding variable modifications to the MaxQuant (deamidation of asparagine and glutamine, oxidation of methionine, acetylation of protein N-term) and Comet (same as MaxQuant plus methylation on lysine, arginine, aspartic and glutamic acid), we observed that the percentage of spliced peptides decreased (Supplemental Figure 3 A, Supplemental Tables 3). The MS/MS scans previously matched to spliced peptides were now matched to modified peptides in UniProt, mainly oxidations (64.7%) and deamidations (28.5%). Again, more conflicting sequences were found in the spliced peptide group (data not shown). If we restricted the analysis to the high quality PSMs, where both MaxQuant and Comet found the same peptide, the overall contribution of *LM_spliced* peptides shrinks further (Supplemental Figure 3 A) to about 2% of the UniProt peptides.

### Synthetic Peptide Searches

We selected 21 MS/MS scans from the Fib sample, where Liepe et al. matched a spliced peptide and MaxQuant matched a UniProt peptide. This selection was not biased to favor MaxQuant results, but we chose spectra that appeared to be typical for the group of spectra that produced conflicting results. For each of the 21 scans we synthesized two peptides: one according to the LM-spliced identification and one for the UniProt alternative. We compared the MS/MS spectra of the synthetic peptides to the spectra of the endogenous peptides. As an example, we show the endogenous and synthetic spectra of the *LM_spliced* peptide DHAQQPYSM (Figure 2 A) and its UniProt competitor peptide DHRSEQSSM (Figure 2 B). All the 21 comparisons can be found in Supplemental Figure 4. Some *LM_spliced* peptides (for examples LENKKGKAL, RVTGALQKK) differ in only two positions from the alternative UniProt peptides (EINKKGKAL, RLSGALQKK), reflecting how similar spliced and UniProt matches can be. We computed the cosine-similarity score between synthetic and endogenous (i.e. original eluted HLA-Ip) spectra. For almost all the 21 cases, synthetic spectra from UniProt peptides fit better to the original MS/MS spectra compared to their *LM_spliced* peptides counterparts (Figure 2 C).

**Figure 2.**
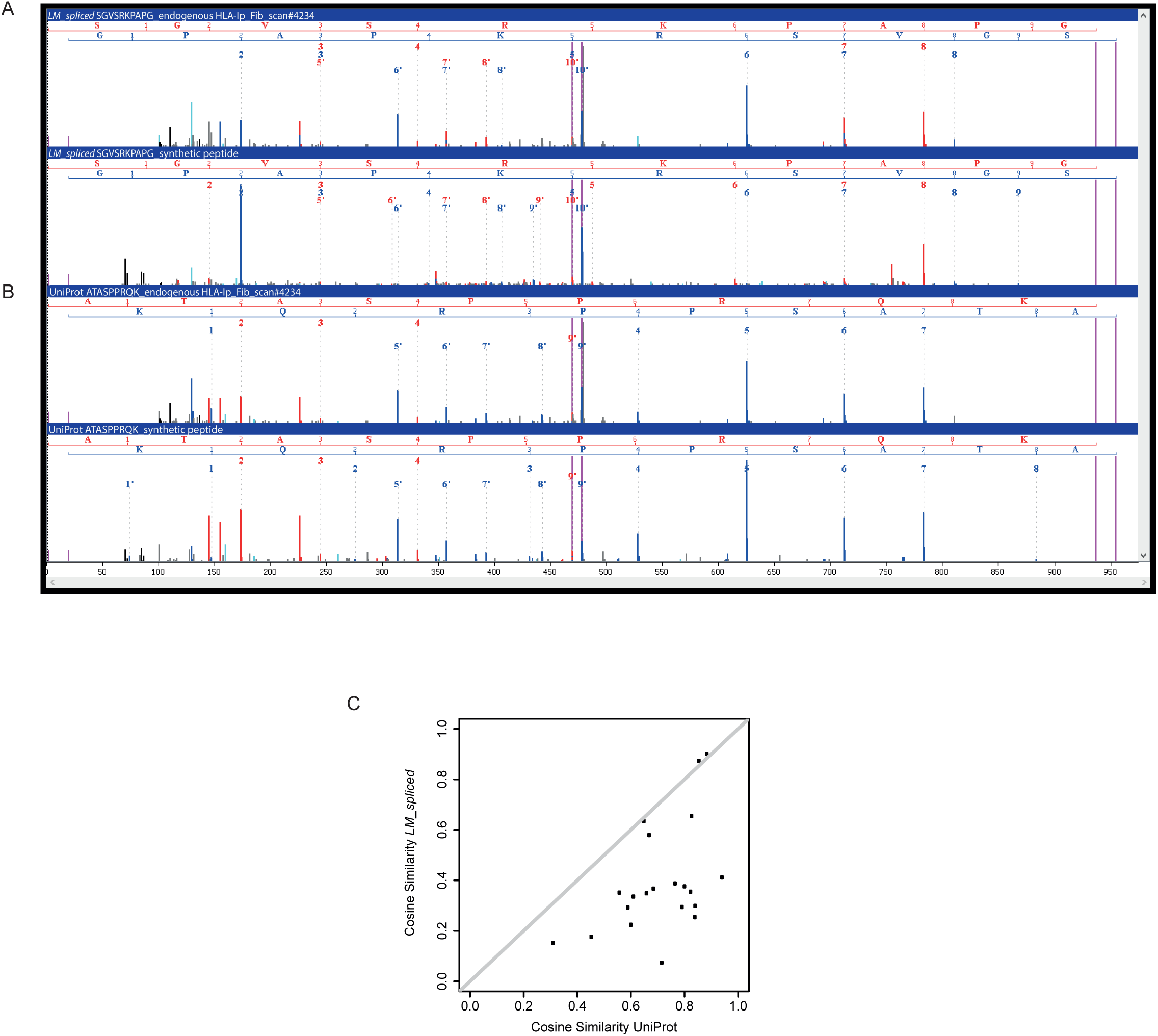
A) Example of MS/MS annotation of endogenous HLA-Ip identified as a *LM_spliced* peptide (DHAQQPYSM), (upper panel) MS/MS of the synthetic counterpart of the *LM_spliced* (lower panel). B) MS/MS annotation of the same endogenous HLA-Ip as an alternative UniProt peptide (DHRSEQSSM, upper panel), and MS/MS of the synthetic counterpart of the alternative UniProt peptide (lower panel). C) The cosine similarity score calculated for the 21 pairs of MS/MS spectra of *LM_spliced* peptides and their synthetic counterparts, and the pairs of the alternative sequences from UniProt and their synthetic peptides.

### Alternative pipeline to estimate the contribution of spliced peptides to the HLA-peptidome

In order to shed more light on the detection of PSPs by MS, we implemented a different computational pipeline (Figure 3 A), which is based on de-novo sequencing (25). This pipeline proceeds in three steps: 1) de-novo sequencing of MS/MS spectra to retrieve only de-novo sequences not found in UniProt. 2) flagging de-novo sequences by the alignment tool TagPep as *DeNovo_spliced* and *DeNovo_non-spliced* peptides. 3) adding the candidate *DeNovo_spliced* and *DeNovo_non-spliced* sequences to UniProt fasta protein/peptide reference database files and searching them with MaxQuant and Comet at FDR of 1% with variable modifications. The idea behind the last step is to match each MS/MS spectrum to either *DeNovo_spliced, DeNovo_non-spliced* or Uniprot. Importantly, this computational pipeline bypasses the step of matching MS/MS spectra to a huge database of potential spliced peptides.

**Figure 3.**
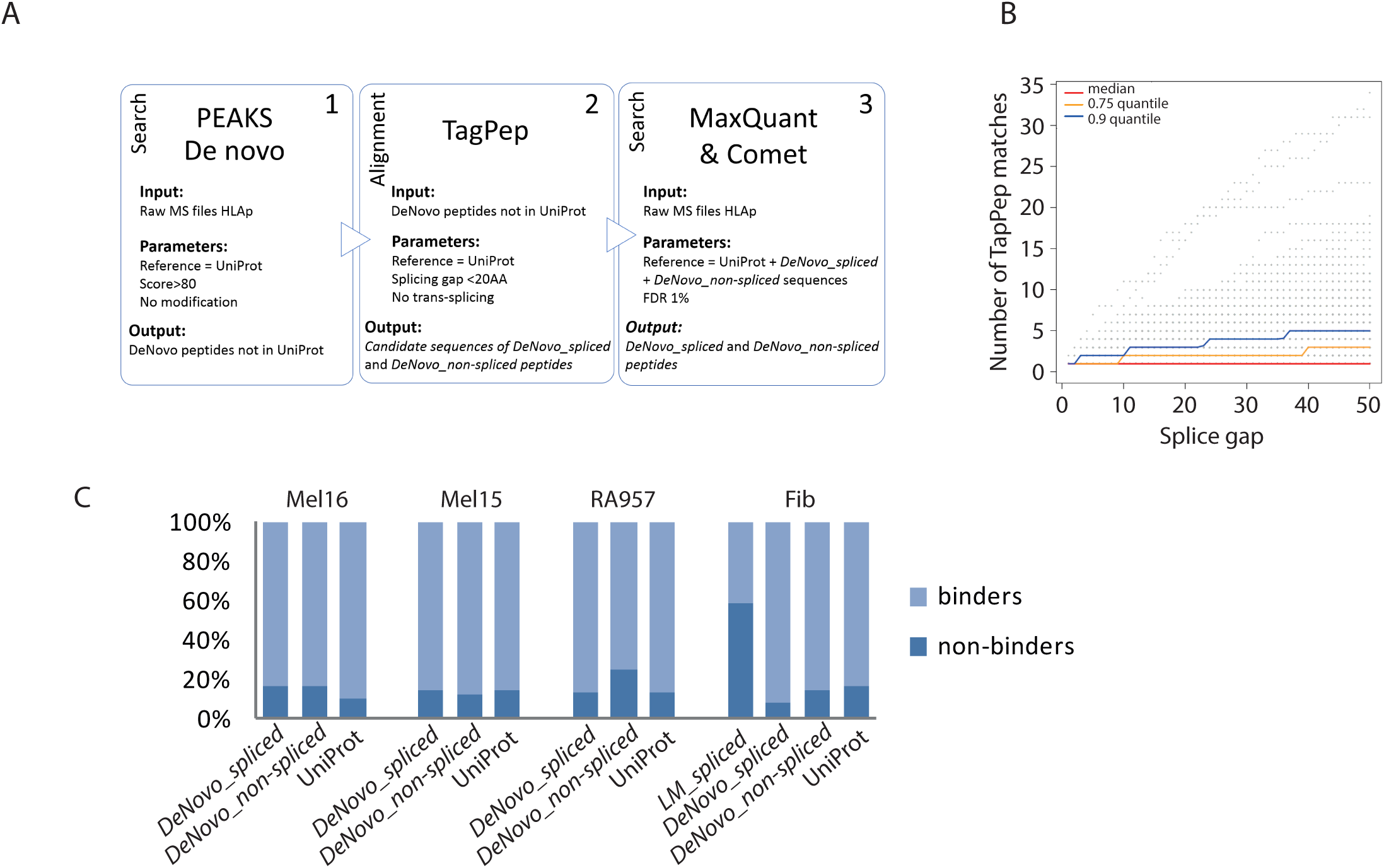
A) Scheme of the de-novo based pipeline to identify possible spliced peptides. B) The splicing gap in relation to the number of TagPep matches. The number of TagPep matches is the number of returned hits with a splicing gap shorter than a given value. Hits with the same splicing position and splicing gap are merged and counted as one hit. C) Distribution of 8-14 mer peptides predicted by NetMHCpan as binders and non-binders among the *DeNovo_spliced, DeNovo_non-spliced* and UniProt peptides identified in Mel16, Mel15, RA957 and Fib samples, and in addition also *LM_spliced* peptides.

We used the PEAKS tool (13) for de-novo sequencing of the immunopeptidomics MS/MS spectra from four different biological samples (see Experimental procedures section for more details). First, we estimated the error present in the PEAKS de-novo sequences. We compared spectra matches assigned by both PEAKS and MaxQuant to UniProt peptides as a function of the PEAKS local confidence score. For a local confidence score higher than 80 the peptides assigned to the spectra by both PEAKS and MaxQuant agreed in 80% of the cases (Supplemental Figure 5 A). We can accept this level of performance, since in our computational approach the proposed peptides are subsequently filtered by the consecutive MaxQuant or Comet analysis at FDR of 1%. Comparison for the UniProt sequences showed that PEAKS could identify about half the peptides as compared to MaxQuant at the given thresholds (Supplemental Figure 5 B). This is expected since de-novo sequencing requires searching a large search space of all AA combinations. Furthermore, we observed that PEAKS is not optimized for non-tryptic peptides (see below).

Next we checked whether the identified de-novo sequences that did not match a UniProt sequence could be PSPs by an in-house alignment tool called TagPep (14). TagPep is a very fast alignment tool employing efficient indexing. TagPep first matches the whole de-novo peptide sequence to the protein database and if there is no hit it tries to match it with one splicing event (Supplemental Data 1). The number of TagPep matches per de-novo sequence depends on the allowed splice gap between the spliced protein fragments. Having no restrictions on the splice gap, 20% of random amino acid sequences of length 8-11 could be matched as a spliced peptide to the human proteome database. Figure 3 B shows that more than half of the *de-novo_spliced* sequences produce a unique TagPep match with a splice gap of less than 20 AA, whereas 90% of the sequences have three matches or less. Only about 10% of the sequences have more than 3 matches and have many possible explanations. Furthermore, we estimated that including the second best PSMs identified by PEAKS de-novo with a local confidence score >80 would increase the number of *de novo sequences by less than 10%* (Supplemental Figure 5 C). For the Mel15 data, TagPep matched 957 peptides as *DeNovo_spliced* when only the best scored PEAKS PSM matches were analyzed (Supplemental Data 1). Including the second best PSM matches resulted in additional 62 TagPep *DeNovo_spliced* matches. Therefore, including lower ranking PEAKS de-novo PSM hits will not drastically affect our results.

Next, for each biological sample we created a separate fasta file by adding the list of *DeNovo_spliced* and the *DeNovo_non-spliced* peptide sequences to the UniProt fasta files. All MS/MS spectra from each immunopeptidomics sample were subsequently matched against these fasta databases using the sequence search tools MaxQuant and Comet. To test the performance of our pipeline we used previously published immunopeptidomics high quality datasets (Mel15, Mel16, Fib and RA957 samples), which represent a variety of HLA binding specificities (Supplemental Table 1). For the Fib, RA957, Mel15 and Mel16 samples we identified 105 (135,84), 140 (137,83), 546 (550,359) and 99 (146,99) spliced peptides with MaxQuant (Comet, consensus of MaxQuant and Comet), respectively. The MaxQuant-Comet consensus *De-Novo spliced* identifications constitutes 1.7%, 1.1%, 2.1% and 0.7% of the identified UniProt immunopeptidome (Supplemental Table 4). Most of the PSPs found by our pipeline were predicted to bind to the HLA-molecules (Figure 3 C). Moreover, in both the PSPs found by our pipeline as well as the *DeNovo_non-spliced*, we could see evidences of the expected binding motifs (Supplemental Figure 6), even if the number of peptides is significantly smaller than for the predicted spliced peptides of Liepe (3). The differences between the binding specificities of *DeNovo_spliced, DeNovo_non-spliced* and UniProt peptides, as seen in case of the HLA-B27:05 in Mel15, are most likely related to a bias against identification of HLA-B27 peptides in PEAKS (Supplemental Figure 7), which has difficulty identifying such non-tryptic peptides.

Compared to the UniProt peptides, the *DeNovo_spliced* peptides found by our pipeline have a slightly higher absolute mass error and lower delta score, but they have very similar score and charge distribution. However, compared to the spliced peptides found by Liepe et al, the *DeNovo_spliced* peptides have better match characteristics: lower absolute mass error, higher Andromeda scores and delta scores, higher number of matching ions and less singly charged PSMs (Supplemental Figure 2 A-F).

### Sequence variants and spliced peptide conflicts

Next, we searched for single amino acids variations (SAAV) obtained by exome sequencing of the Mel15 and Mel16 samples (10). The spliced peptides found by MaxQuant for the Mel15 and Mel 16 samples were added to the fasta file used by Bassani-Sternberg et al., which contained all *Ensemble* human protein sequences and the sequence variants identified by exome sequencing. Out of the 600 *DeNovo_spliced* peptides, 17 (2.8%) unique peptides had the same sequence as an endogenous HLA-Ip with a single amino acid variation (Supplemental Data 13). Ten of the *DeNovo_non-spliced* sequences, which were identified by our pipeline, could also be explained by a peptide encompassing such variants. These results highlight the needs to carefully evaluate spliced peptides identified by MS/MS and make sure that they do not have a different, potentially simpler explanation.

### Characterization of the splicing events

Lastly, we found that in many *DeNovo_spliced* peptides, identified by both MaxQuant and Comet, the splicing position is at the N- and C-termini (Figure 4 A) in contrast to the *LM_spliced* peptides (Figure 4 B), which have more uniformly distributed splicing positions. The distribution of MaxQuant delta scores as a function of splice position indicates that certain splice positions may on average be more ambiguous than others (Figure 4 C). On the other hand, it could also be possible that single AA abundant in the proteasome or during sample processing could be attached to the peptide termini. This effect is known as transpeptidation and was observed in tryptic samples (26).

**Figure 4.**
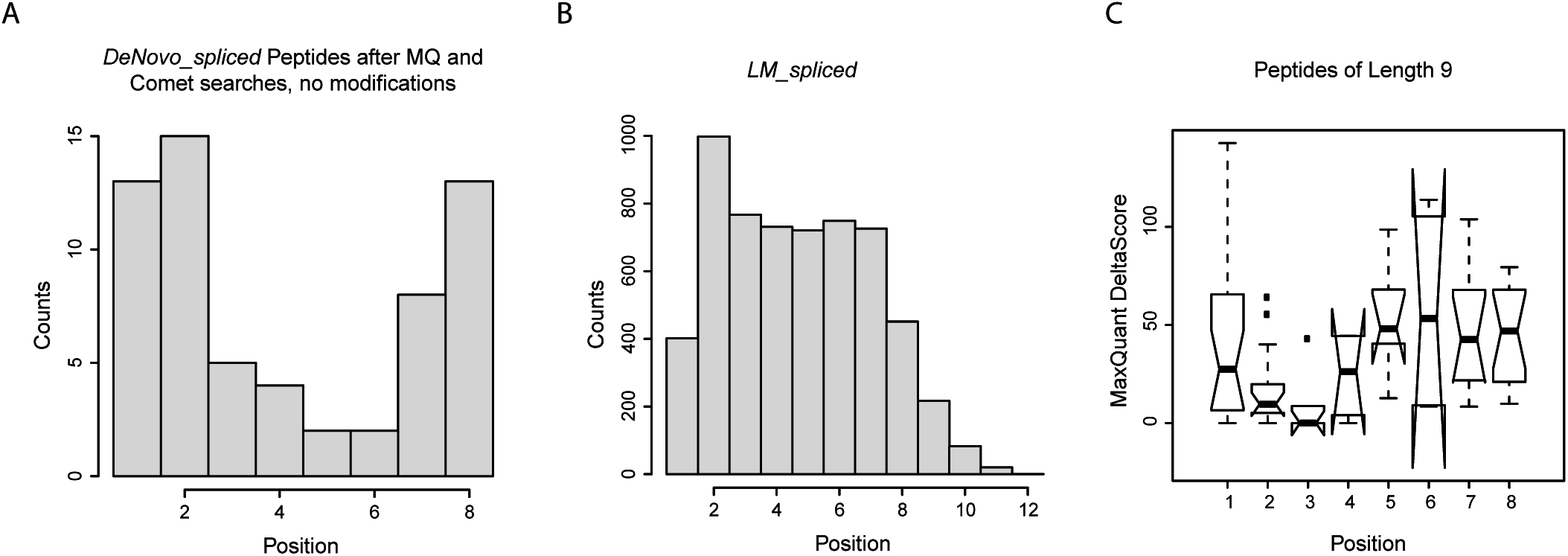
A) Histogram of splicing positions within 9-mers for the *DeNovo_spliced* peptides identified by both Comet and MaxQuant. B) Histogram of *LM_spliced* peptides splicing positions. C) MaxQuant delta scores as a function of the splicing position within the – 9-mers for the *DeNovo_spliced* peptides identified by both Comet and MaxQuant.

## Discussion

Examples of proteasomal-spliced peptides have been reported and some of them were shown to be immunogenic (4-9, 16, 27-29). A true estimation of the contribution of spliced peptides to the global immunopeptidome is critical in order to fundamentally understand the biological pathways involved. Consequently, advanced computational and experimental tools must be developed and benchmarked to facilitate their confident identification.

Liepe et al. (3) were the first to attempt to find PSPs by means of MS on a large scale. Their report concluded that about 30% of the HLA-Ip are produced by proteasomal splicing by transpeptidation of two noncontiguous fragments of a parental protein (cis-splicing). Our reanalysis of their results revealed that their approach led to the identification of PSPs candidates that did not fit the consensus binding motifs, while the non-spliced UniProt HLA-Ip identified in the same experiment did. Additional parameters related to their MS/MS spectrum matches suggest that many of the spliced peptide matches reported in Liepe et al. are ambiguous and were ruled out when we used different search engines and included common PTMs or sequence variants in the search. We postulate that because of the huge search space of potential spliced peptide, the bioinformatics approach applied by Liepe et al. led to uncontrollable propagation of false positives. The effect of database size and the increased likelihood of false positive identifications in proteogenomics applications have been thoroughly reviewed in (30), and these concepts are relevant here as well. Therefore, the true contribution of spliced peptides to the immunopeptidome has yet to be defined.

In a typical peptidomics setting, we match MS/MS spectra against a large set of theoretical peptide spectra, most of which are not present in the sample. This endeavor produces two types of PSMs: true matches and false positives. False positives are very common especially for spliced peptides since these peptides can produce similar MS/MS spectra to UniProt peptides with similar match scores. For example, the potential spliced peptide KRI-PLPTKK only differs from its UniProt counterpart RIKPLPTKK by a permutation of the first three AA. When using a very large proteasomal spliced peptide database there is an elevated chance that a potential spliced peptide will have a very similar spectrum to the UniProt peptide and produce a higher match score. Furthermore, if a spectrum has no match in the UniProt database (e.g. when it originates from a modified peptide, sequence variant or contaminant not considered in the search) it may still match a spliced peptide with a score that is significant.

The error in the multiple testing of MS/MS searches is controlled by using decoy database in order to calculate the FDR (31, 32). One assumption behind this target-decoy approach is that the scores of the decoy peptides reflect the scores of wrongly assigned PSMs. When using decoys for spliced peptides, their similarity with the UniProt sequences may be lost and one would have to carefully evaluate whether the assumption mentioned above still holds. If it does not hold, the target-decoy approach might underestimate the FDR and lead to many false positives especially for large spliced peptide databases.

Trans-splicing of fragments derived from two source proteins that happen to be present in the same proteasome complex at the same time, is unlikely to happen on a large scale, hence we focused our study on cis-splicing events. To overcome biases related to searching all possible cis-splice peptides, we developed an alternative workflow based on de-novo sequencing and subsequent verification with multiple search tools including the most prevalent amino acid modifications and sequence variants detected by exome sequencing. We found that 1-2% of the high quality PSMs originate from potential proteasome cis-spliced peptides. These peptides fitted the HLA consensus binding motifs and had good spectrum match properties. Given that our de-novo sequencing approach finds about half of the peptides compared to a UniProt sequence search, we can say that the maximal amount of spliced peptide candidates is 2-4%. However, MS/MS based approaches cannot ultimately determine the creation mechanism of these peptides and different sequence interpretations may also be possible. For example, a significant number of detected HLA-Ip originates from transcripts, which do not fall into a UniProt protein coding region (33), and these non-canonical peptides could be misinterpreted as PSPs. Other ambiguities may be due to post-translational or chemical peptide modifications not considered in the search. Overall, our results present an upper bound for the proportion of cis-spliced peptides, and the true contribution of such PSPs to the HLA-I ligandome may be much smaller.

## Footnotes

We are thankful to Peter A. van Veelen for critically reading our manuscript. This work was supported by the Ludwig Institute for Cancer Research and by the ISREC Foundation thanks to a donation from the Biltema Foundation.

## References

1. Neefjes, J., Jongsma, M. L., Paul, P., and Bakke, O. (2011) Towards a systems understanding of MHC class I and MHC class II antigen presentation. Nature reviews. Immunology 11, 823–836

2. Bassani-Sternberg, M., Pletscher-Frankild, S., Jensen, L. J., and Mann, M. (2015) Mass spectrometry of human leukocyte antigen class I peptidomes reveals strong effects of protein abundance and turnover on antigen presentation. Molecular & cellular proteomics : MCP 14, 658–673

3. Liepe, J., Marino, F., Sidney, J., Jeko, A., Bunting, D. E., Sette, A., Kloetzel, P. M., Stumpf, M. P., Heck, A. J., and Mishto, M. (2016) A large fraction of HLA class I ligands are proteasome-generated spliced peptides. Science 354, 354–358

4. Ebstein, F., Textoris-Taube, K., Keller, C., Golnik, R., Vigneron, N., Van den Eynde, B. J., Schuler-Thurner, B., Schadendorf, D., Lorenz, F. K. M., Uckert, W., Urban, S., Lehmann, A., Albrecht-Koepke, N., Janek, K., Henklein, P., Niewienda, A., Kloetzel, P. M., and Mishto, M. (2016) Proteasomes generate spliced epitopes by two different mechanisms and as efficiently as non-spliced epitopes. 6, 24032

5. Hanada, K., Yewdell, J. W., and Yang, J. C. (2004) Immune recognition of a human renal cancer antigen through post-translational protein splicing. Nature 427, 252–256

6. Vigneron, N., Stroobant, V., Chapiro, J., Ooms, A., Degiovanni, G., Morel, S., van der Bruggen, P., Boon, T., and Van den Eynde, B. J. (2004) An antigenic peptide produced by peptide splicing in the proteasome. Science 304, 587–590

7. Warren, E. H., Vigneron, N. J., Gavin, M. A., Coulie, P. G., Stroobant, V., Dalet, A., Tykodi, S. S., Xuereb, S. M., Mito, J. K., Riddell, S. R., and Van den Eynde, B. J. (2006) An antigen produced by splicing of noncontiguous peptides in the reverse order. Science 313, 1444–1447

8. Dalet, A., Robbins, P. F., Stroobant, V., Vigneron, N., Li, Y. F., El-Gamil, M., Hanada, K., Yang, J. C., Rosenberg, S. A., and Van den Eynde, B. J. (2011) An antigenic peptide produced by reverse splicing and double asparagine deamidation. Proceedings of the National Academy of Sciences of the United States of America 108, E323–331

9. Michaux, A., Larrieu, P., Stroobant, V., Fonteneau, J. F., Jotereau, F., Van den Eynde, B. J., Moreau-Aubry, A., and Vigneron, N. (2014) A spliced antigenic peptide comprising a single spliced amino acid is produced in the proteasome by reverse splicing of a longer peptide fragment followed by trimming. Journal of immunology 192, 1962–1971

10. Bassani-Sternberg, M., Braunlein, E., Klar, R., Engleitner, T., Sinitcyn, P., Audehm, S., Straub, M., Weber, J., Slotta-Huspenina, J., Specht, K., Martignoni, M. E., Werner, A., Hein, R. D H. B., Peschel, C., Rad, R., Cox, J., Mann, M., and Krackhardt, A. M. (2016) Direct identification of clinically relevant neoepitopes presented on native human melanoma tissue by mass spectrometry. Nature communications 7, 13404

11. Bassani-Sternberg, M., Chong, C., Guillaume, P., Solleder, M., Pak, H., Gannon, P. O., Kandalaft, L. E., Coukos, G., and Gfeller, D. (2017) Deciphering HLA-I motifs across HLA peptidomes improves neoantigen predictions and identifies allostery regulating HLA specificity. PLoS computational biology 13, e1005725

12. Vizcaino, J. A., Cote, R. G., Csordas, A., Dianes, J. A., Fabregat, A., Foster, J. M., Griss, J., Alpi, E., Birim, M., Contell, J., O'Kelly, G., Schoenegger, A., Ovelleiro, D., Perez-Riverol, Y., Reisinger, F., Rios, D., Wang, R., and Hermjakob, H. (2013) The PRoteomics IDEntifications (PRIDE) database and associated tools: status in 2013. Nucleic acids research 41, D1063–1069

13. Ma, B., Zhang, K., Hendrie, C., Liang, C., Li, M., Doherty-Kirby, A., and Lajoie, G. (2003) PEAKS: powerful software for peptide de novo sequencing by tandem mass spectrometry. Rapid Commun Mass Spectrom 17, 2337–2342

14. Iseli, C., Ambrosini, G., Bucher, P., and Jongeneel, C. V. (2007) Indexing strategies for rapid searches of short words in genome sequences. PloS one 2, e579

15. Mishto, M., and Liepe, J. (2017) Post-Translational Peptide Splicing and T Cell Responses. Trends Immunol 38, 904–915

16. Berkers, C. R., de Jong, A., Schuurman, K. G., Linnemann, C., Meiring, H. D., Janssen, L., Neefjes, J. J., Schumacher, T. N., Rodenko, B., and Ovaa, H. (2015) Definition of Proteasomal Peptide Splicing Rules for High-Efficiency Spliced Peptide Presentation by MHC Class I Molecules. Journal of immunology 195, 4085–4095

17. Cox, J., and Mann, M. (2008) MaxQuant enables high peptide identification rates, individualized p.p.b.-range mass accuracies and proteome-wide protein quantification. Nature biotechnology 26, 1367–1372

18. Eng, J. K., Jahan, T. A., and Hoopmann, M. R. (2013) Comet: an open-source MS/MS sequence database search tool. Proteomics 13, 22–24

19. Stein, S. E., and Scott, D. R. (1994) Optimization and testing of mass spectral library search algorithms for compound identification. Journal of the American Society for Mass Spectrometry 5, 859–866

20. Horlacher, O., Nikitin, F., Alocci, D., Mariethoz, J., Muller, M., and Lisacek, F. (2015) MzJava: An open source library for mass spectrometry data processing. Journal of proteomics 129, 63–70

21. Nielsen, M., and Andreatta, M. (2016) NetMHCpan-3.0; improved prediction of binding to MHC class I molecules integrating information from multiple receptor and peptide length datasets. Genome medicine 8, 33

22. Andreatta, M., Alvarez, B., and Nielsen, M. (2017) GibbsCluster: unsupervised clustering and alignment of peptide sequences. Nucleic acids research

23. Andreatta, M., Lund, O., and Nielsen, M. (2013) Simultaneous alignment and clustering of peptide data using a Gibbs sampling approach. Bioinformatics 29, 8–14

24. Bassani-Sternberg, M., and Gfeller, D. (2016) Unsupervised HLA Peptidome Deconvolution Improves Ligand Prediction Accuracy and Predicts Cooperative Effects in Peptide-HLA Interactions. Journal of immunology 197, 2492–2499

25. Ma, B., and Johnson, R. (2012) De novo sequencing and homology searching. Molecular & cellular proteomics : MCP 11, O111 014902

26. Schaefer, H., Chamrad, D. C., Marcus, K., Reidegeld, K. A., Bluggel, M., and Meyer, H. E. (2005) Tryptic transpeptidation products observed in proteome analysis by liquid chromatography-tandem mass spectrometry. Proteomics 5, 846–852

27. Berkers, C. R., de Jong, A., Schuurman, K. G., Linnemann, C., Geenevasen, J. A., Schumacher, T. N., Rodenko, B., and Ovaa, H. (2015) Peptide Splicing in the Proteasome Creates a Novel Type of Antigen with an Isopeptide Linkage. Journal of immunology 195, 4075–4084

28. Dalet, A., Vigneron, N., Stroobant, V., Hanada, K., and Van den Eynde, B. J. (2010) Splicing of distant peptide fragments occurs in the proteasome by transpeptidation and produces the spliced antigenic peptide derived from fibroblast growth factor-5. Journal of immunology 184, 3016–3024

29. Platteel, A. C. M., Liepe, J., Textoris-Taube, K., Keller, C., Henklein, P., Schalkwijk, H. H., Cardoso, R., Kloetzel, P. M., Mishto, M., and Sijts, A. (2017) Multi-level Strategy for Identifying Proteasome-Catalyzed Spliced Epitopes Targeted by CD8(+) T Cells during Bacterial Infection. Cell reports 20, 1242–1253

30. Nesvizhskii, A. I. (2014) Proteogenomics: concepts, applications and computational strategies. Nature methods 11, 1114–1125

31. Benjamini, Y., and Hochberg, Y. (1995) Controlling the False Discovery Rate: A Practical and Powerful Approach to Multiple Testing. Journal of the Royal Statistical Society. Series B (Methodological) 57, 289–300

32. Choi, H., and Nesvizhskii, A. I. (2008) False discovery rates and related statistical concepts in mass spectrometry-based proteomics. Journal of proteome research 7, 47–50

33. Pearson, H., Daouda, T., Granados, D. P., Durette, C., Bonneil, E., Courcelles, M., Rodenbrock, A., Laverdure, J. P., Cote, C., Mader, S., Lemieux, S., Thibault, P., and Perreault, C. (2016) MHC class I-associated peptides derive from selective regions of the human genome. The Journal of clinical investigation 126, 4690–4701

