## Supplementary Materials for "Estimating the Contribution of Proteasomal Spliced Peptides to the HLA-I Ligandome" (from another version of this preprint: version 1, 2018-03-26)

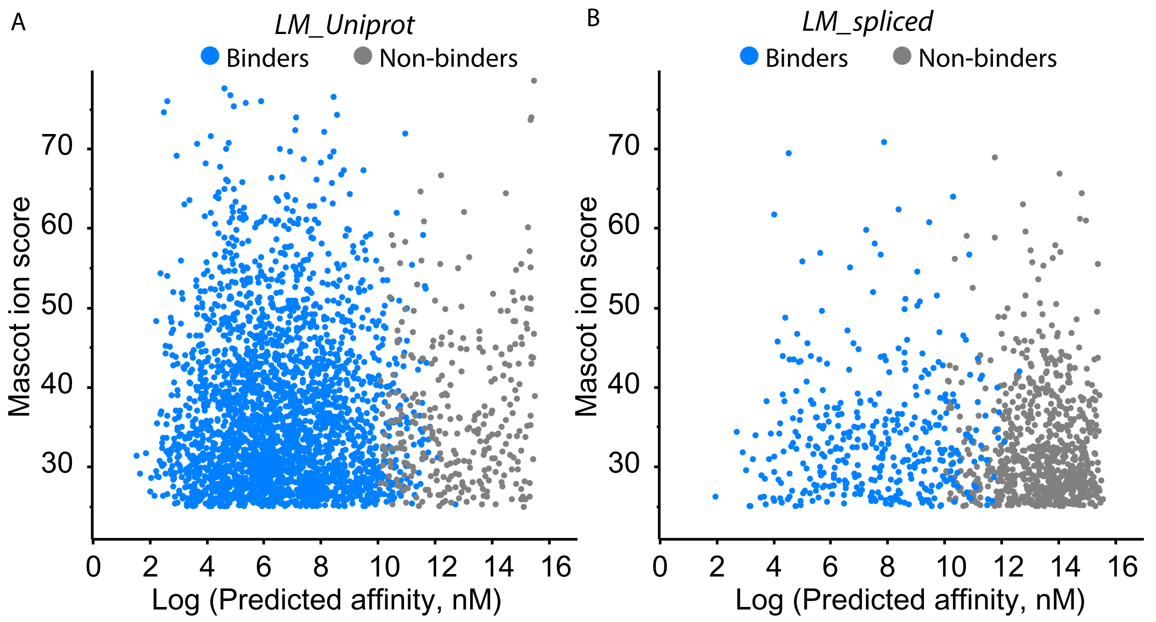


Supplementary Figure 1: Re-analysis of the Liepe et al. data available as Supplementary Data 5. A) Correlation of the predicted binding affinity by NetMHCpan and MS/MS identification scores of peptides identified as LM_UniProt (A) and LM_spliced (B). Binders (rank top 2%) are marked in blue and non-binders in gray.

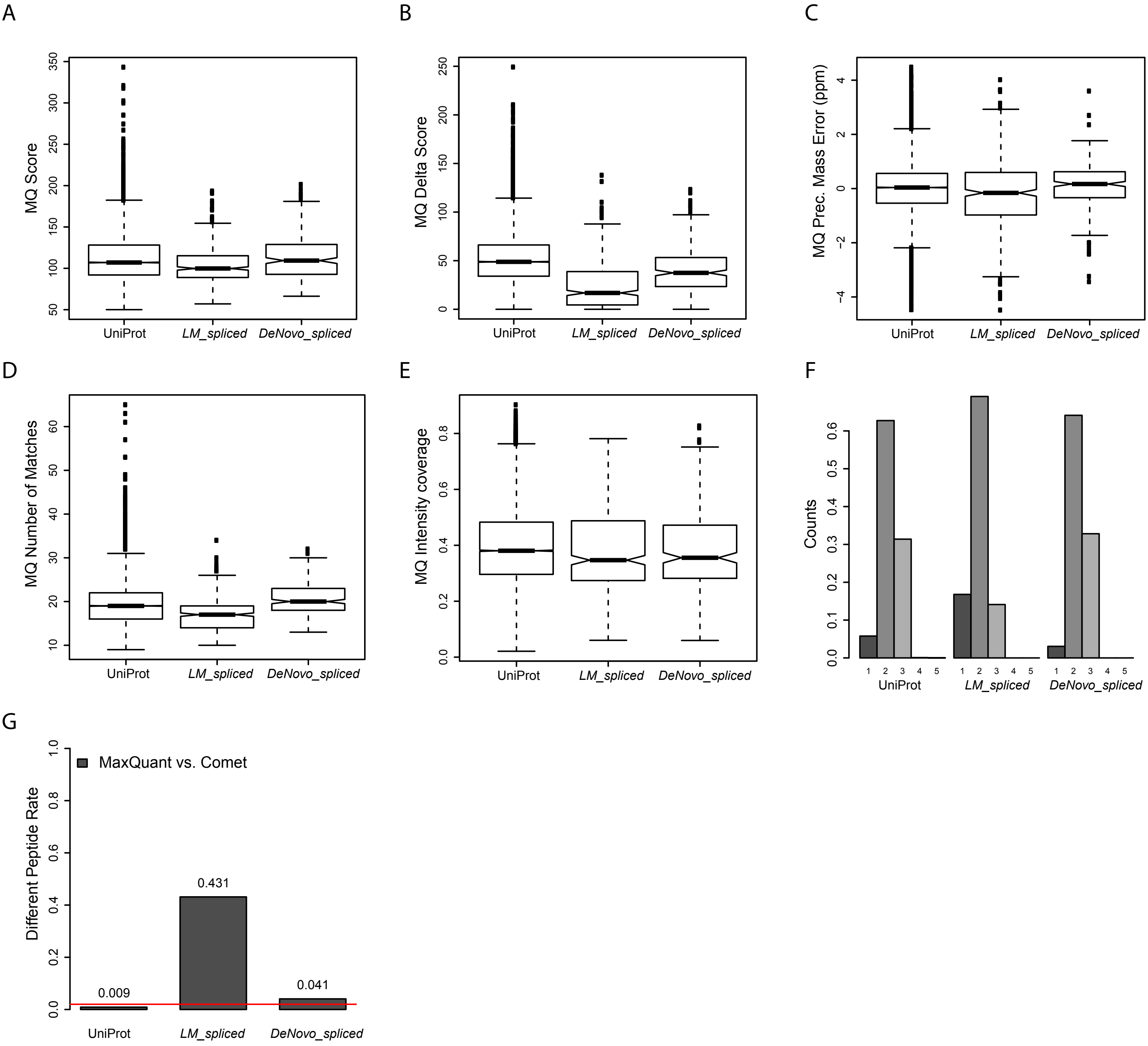


Supplementary Figure 2: Compared to UniProt and *DeNovo_spliced* peptides, *LM_spliced* peptides were characterized with A) lower Andromeda score for the best MS/MS spectrum, B) lower score difference to the second best identified peptide, C) higher absolute precursor mass deviation, D) fewer peaks matching to the predicted fragmentation spectrum, E) lower fraction of total MS/MS peak intensity matched, and F) a larger fraction of singly charged MS/MS spectra matched. G) Rate of disagreement in peptide identification between MaxQuant and Comet for MS/MS scans matched to Uniprot, *LM_spliced* and *DeNovo_spliced* peptides.

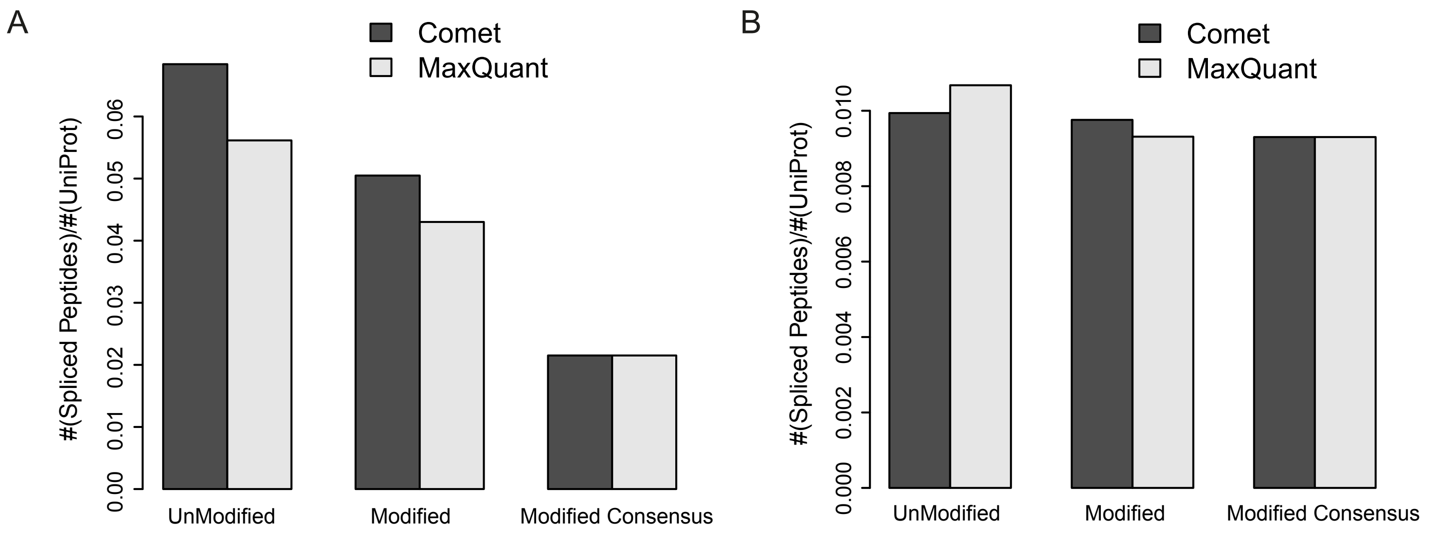


Supplementary Figure 3: Ratio of spliced peptide count and UniProt peptide count for the *LM_spliced* (A) and *DeNovo_spliced* (B) peptides for all Fib samples. Variable modifications are as specified in the text. For 'Modified consensus' the peptide counts are calculated on the subset of PSMs where Comet and MaxQuant agreed, therefore, the values for Comet and MaxQuant are the same.

Supplementary Figure 4: Comparison of MS/MS annotations of endogenous and synthetic counterparts of 21 pairs of *LM_spliced* and their UniProt alternatives, and of the three *LM_spliced* peptides. For the 21 pairs we provide: MS/MS annotation of endogenous HLA-Ip as *LM_spliced* peptides, MS/MS annotation of the synthetic counterpart of the *LM_spliced*, MS/MS annotation of the same endogenous HLA-Ip as an alternative UniPro*t* peptide, and MS/MS annotation of the synthetic counterpart of the alternative UniProt peptide. For the three *LM_spliced* peptides we provide MS/MS annotations of endogenous HLA-Ip and of the synthetic counterparts.

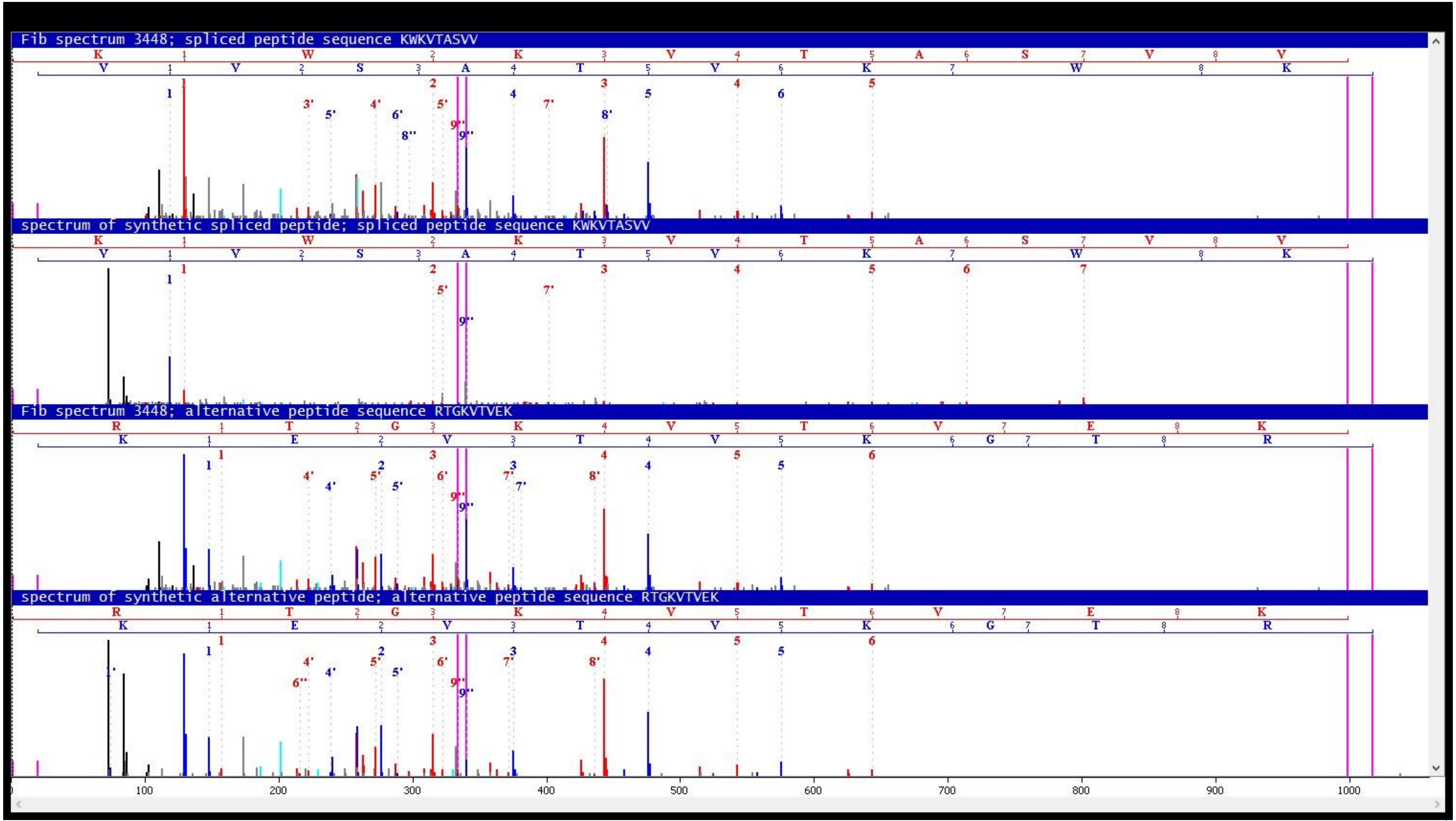
Fib spectrum 3448; spliced peptide sequence KWKVTASVV
spectrum of synthetic spliced peptide; spliced peptide sequence KWKVTASVV
Fib spectrum 3448; alternative peptide sequence RTGKVTVEK
spectrum of synthetic alternative peptide; alternative peptide sequence RTGKVTVEK
100
200
300
400
500
600
700
800
900
1000

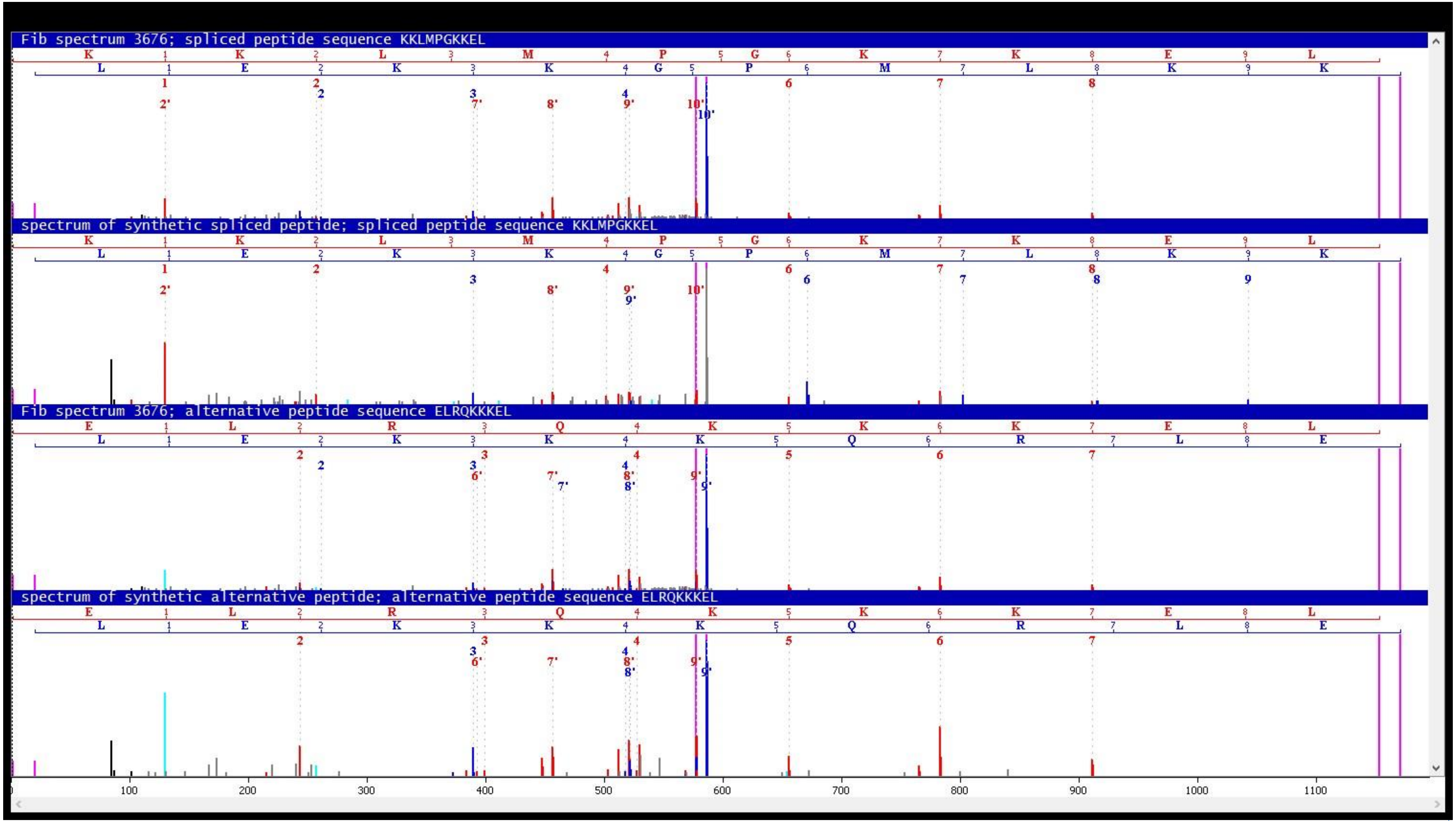
Fib spectrum 3676; spliced peptide sequence KKLMPGKKEL
spectrum of synthetic spliced peptide; spliced peptide sequence KKLMPGKKEL
Fib spectrum 3676; alternative peptide sequence ELRQKKKEL
spectrum of synthetic alternative peptide; alternative peptide sequence ELRQKKKEL
100
200
300
400
500
600
700
800
900
1000
1100

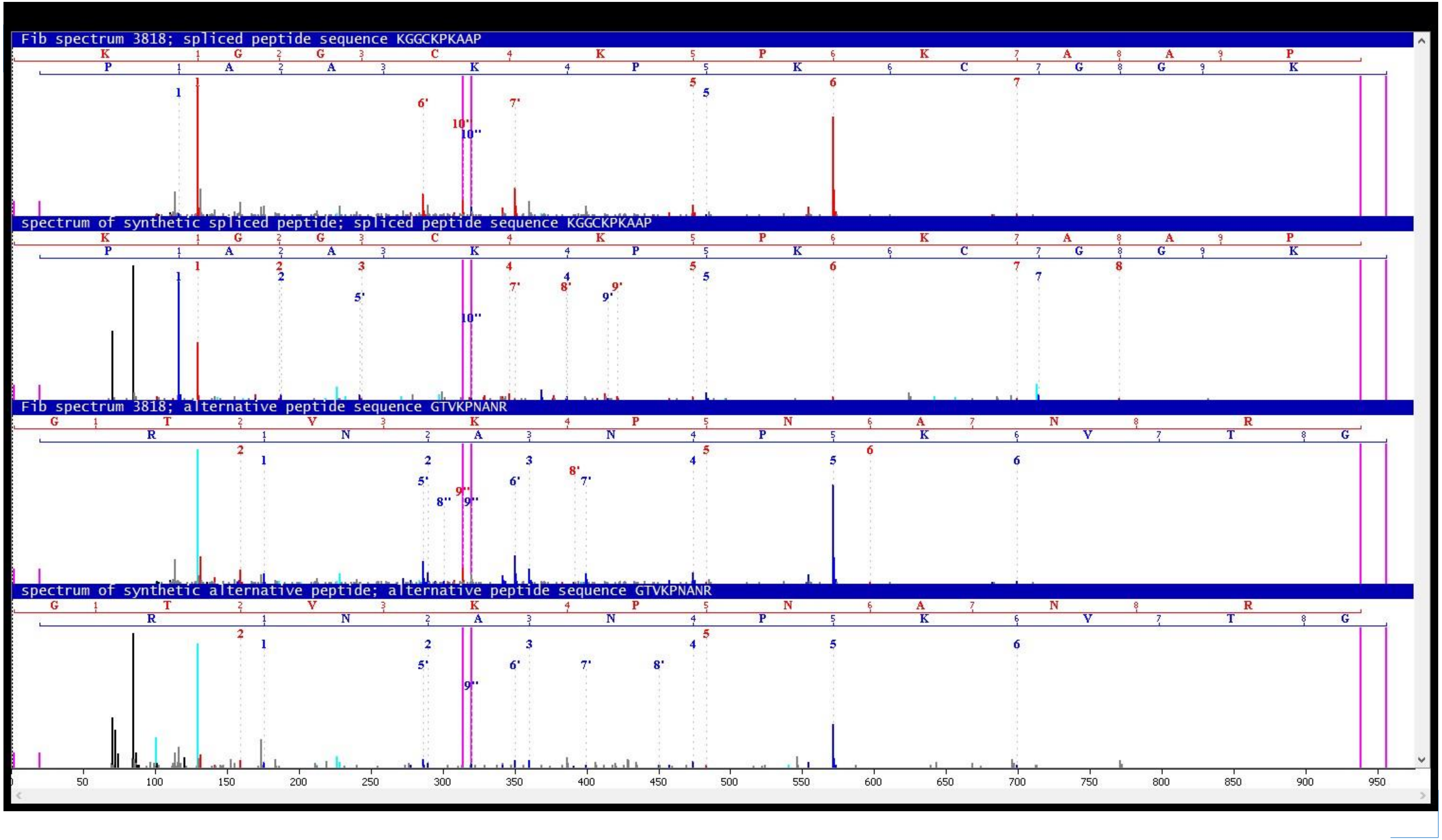
Fib spectrum 3818; spliced peptide sequence KGGCKPKAAP
spectrum of synthetic spliced peptide; spliced peptide sequence KGGCKPKAAP
Fib spectrum 3818; alternative peptide sequence GTVKPNANR
spectrum of synthetic alternative peptide; alternative peptide sequence GTVKPNANR
50 100 150 200 250 300 350 400 450 500 550 600 650 700 750 800 850 900 950

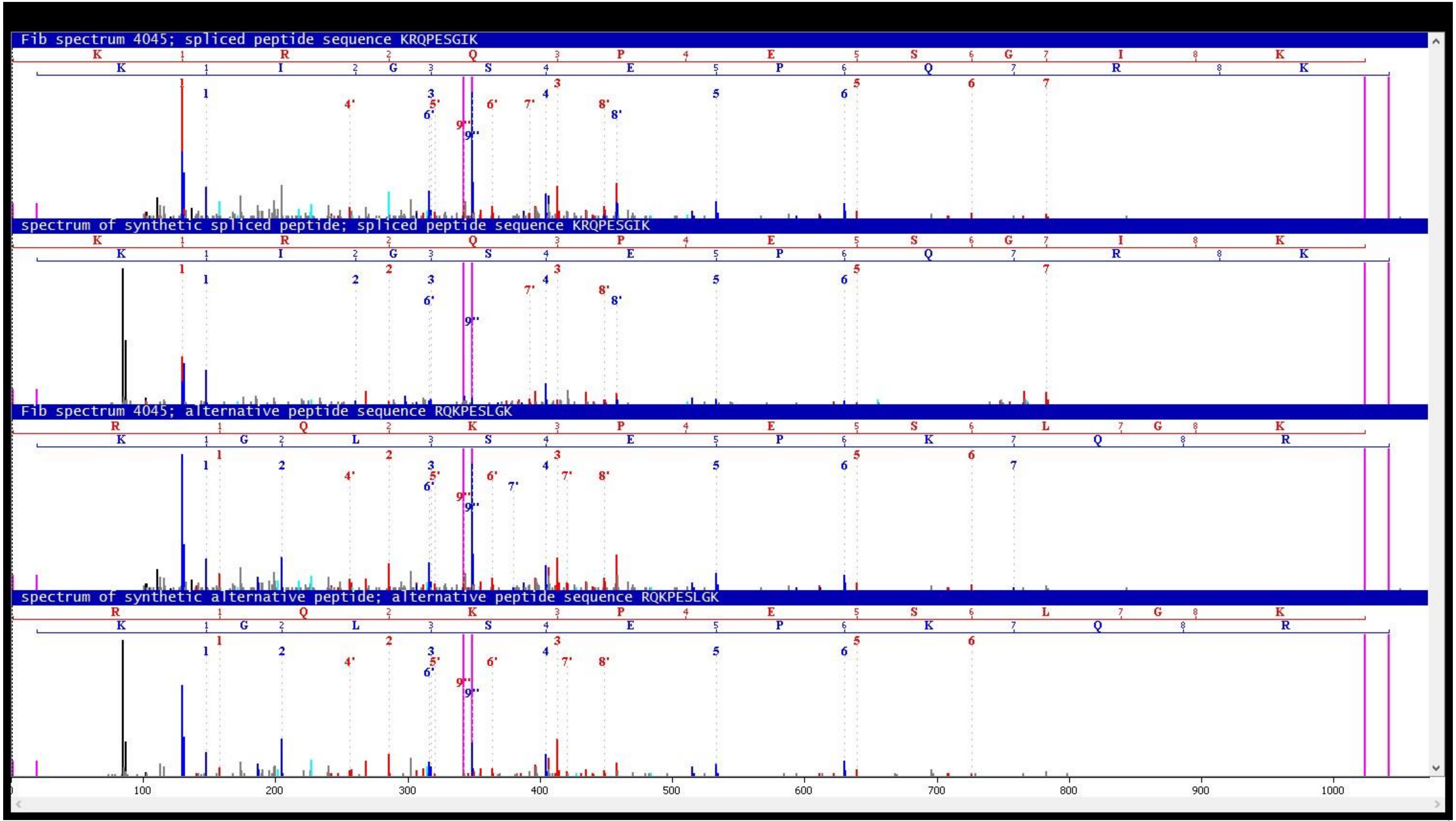

Fib spectrum 4045; spliced peptide sequence KRQPESGIK
spectrum of synthetic spliced peptide; spliced peptide sequence KRQPESGIK
Fib spectrum 4045; alternative peptide sequence RQKPESLGK
spectrum of synthetic alternative peptide; alternative peptide sequence RQKPESLGK
100
200
300
400
500
600
700
800
900
1000

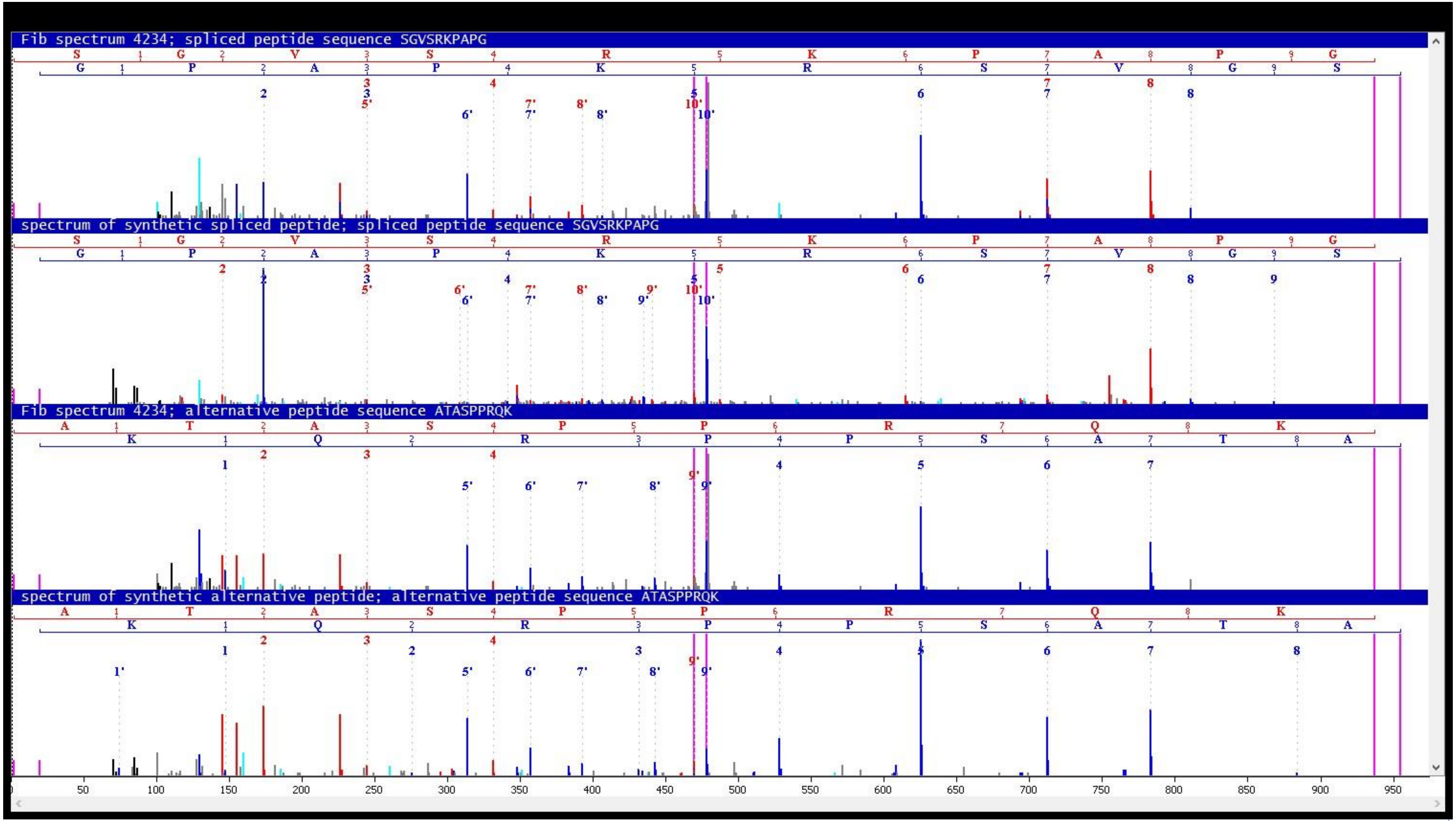
Fib spectrum 4234; spliced peptide sequence SGVSRKPAPG
spectrum of synthetic spliced peptide; spliced peptide sequence SGVSRKPAPG
Fib spectrum 4234; alternative peptide sequence ATASPPRQK
spectrum of synthetic alternative peptide; alternative peptide sequence ATASPPRQK
50 100 150 200 250 300 350 400 450 500 550 600 650 700 750 800 850 900 950

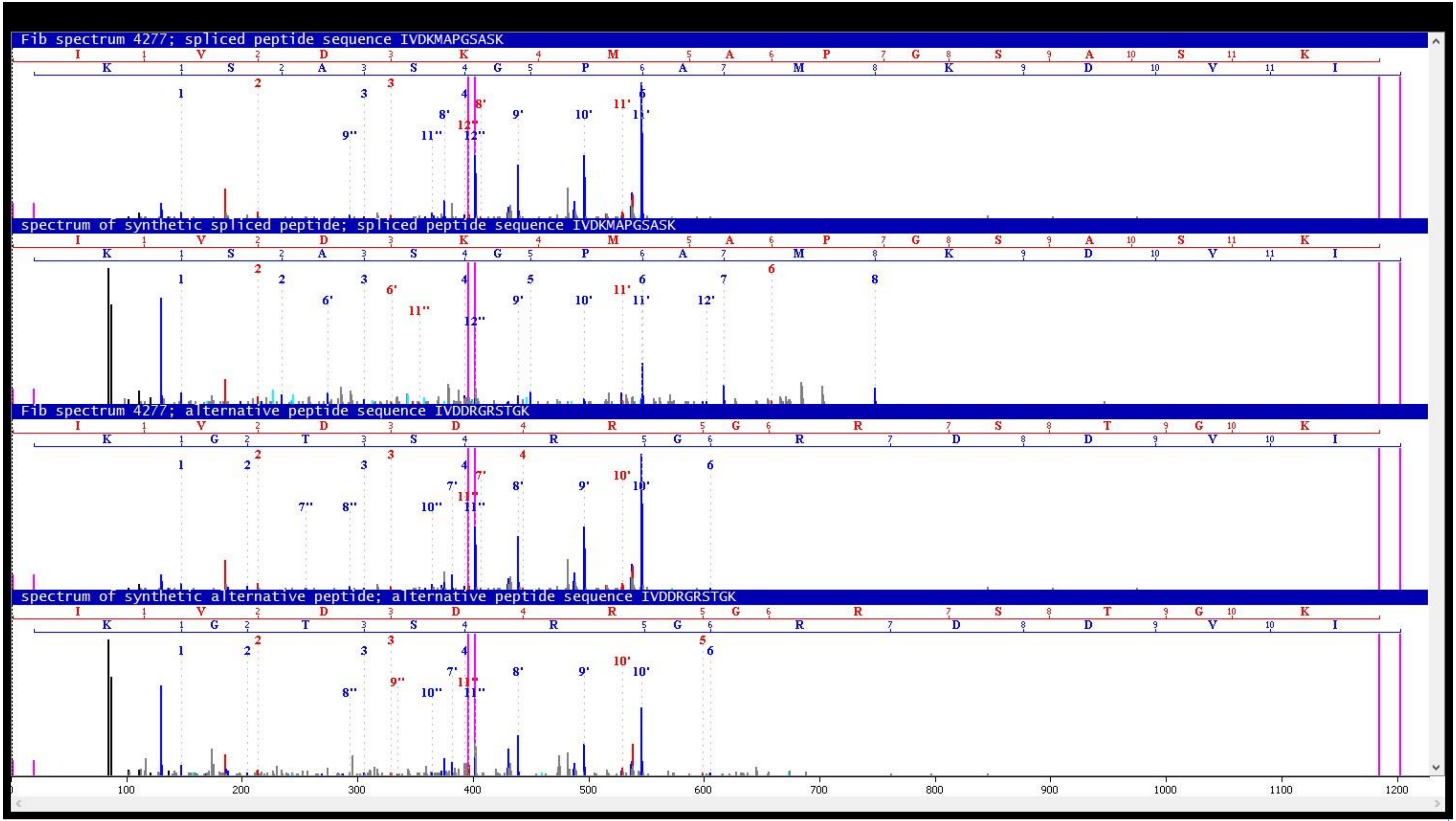
Fib spectrum 4277; spliced peptide sequence IVDKMAPGSASK
spectrum of synthetic spliced peptide; spliced peptide sequence IVDKMAPGSASK
Fib spectrum 4277; alternative peptide sequence IVDDRGRSTGK
spectrum of synthetic alternative peptide; alternative peptide sequence IVDDRGRSTGK
100
200
300
400
500
600
700
800
900
1000
1100
1200

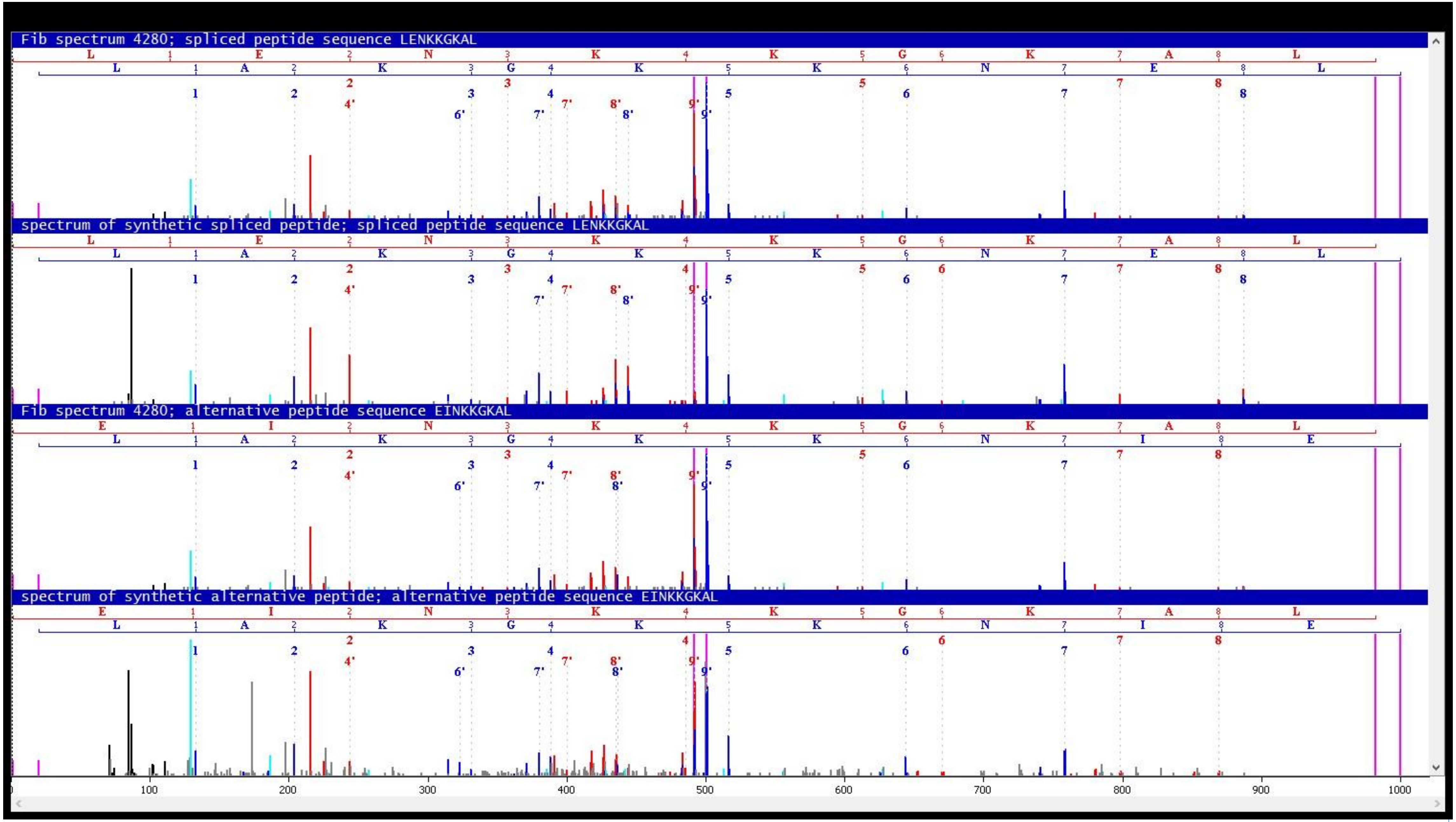

Fib spectrum 4280; spliced peptide sequence LENKKGKAL
spectrum of synthetic spliced peptide; spliced peptide sequence LENKKGKAL
Fib spectrum 4280; alternative peptide sequence EINKKGKAL
spectrum of synthetic alternative peptide; alternative peptide sequence EINKKGKAL
100
200
300
400
500
600
700
800
900
1000

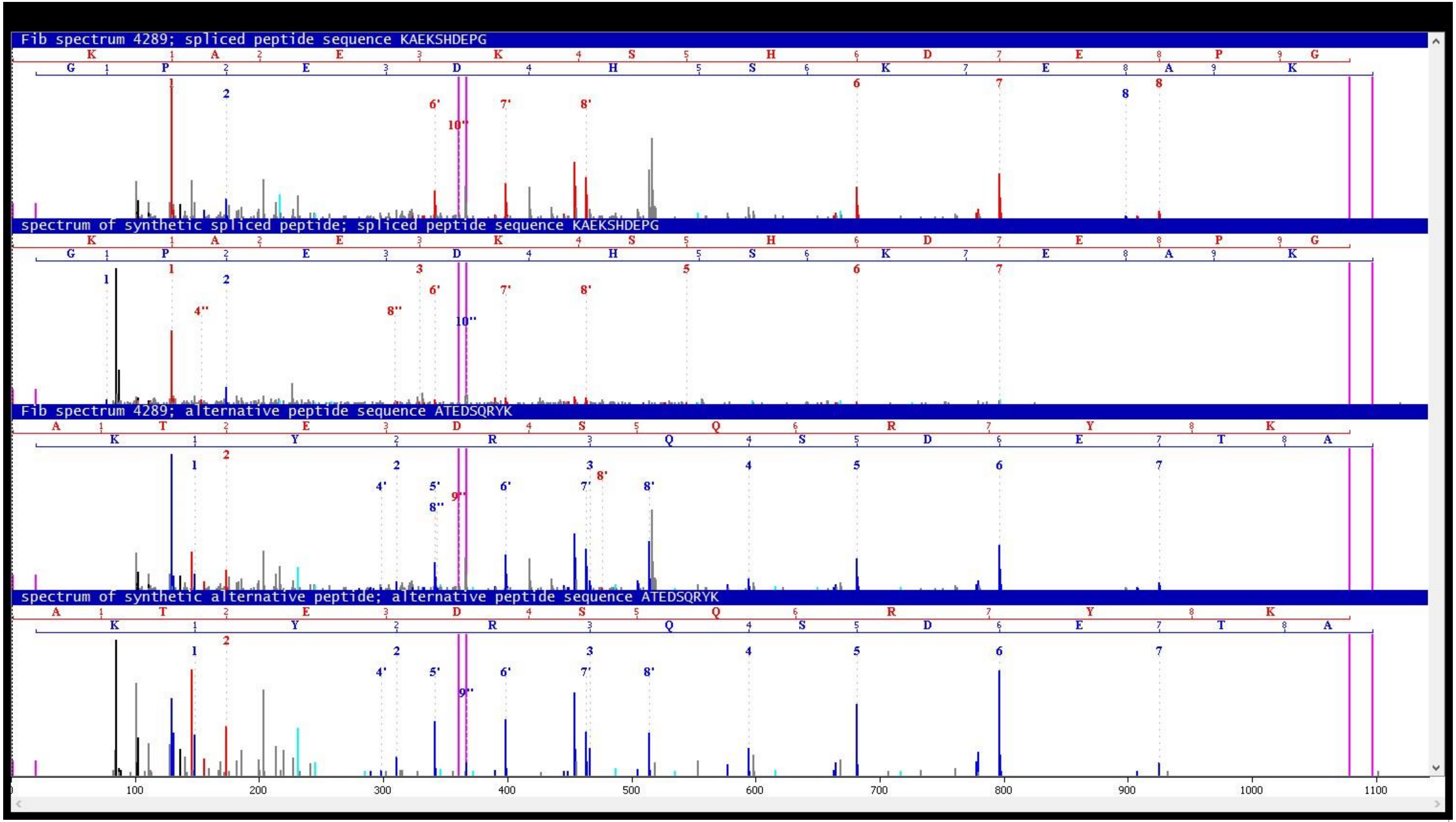
Fib spectrum 4289; spliced peptide sequence KAEKSHDEPG
spectrum of synthetic spliced peptide; spliced peptide sequence KAEKSHDEPG
Fib spectrum 4289; alternative peptide sequence ATEDSQRYK
spectrum of synthetic alternative peptide; alternative peptide sequence ATEDSQRYK
100
200
300
400
500
600
700
800
900
1000
1100

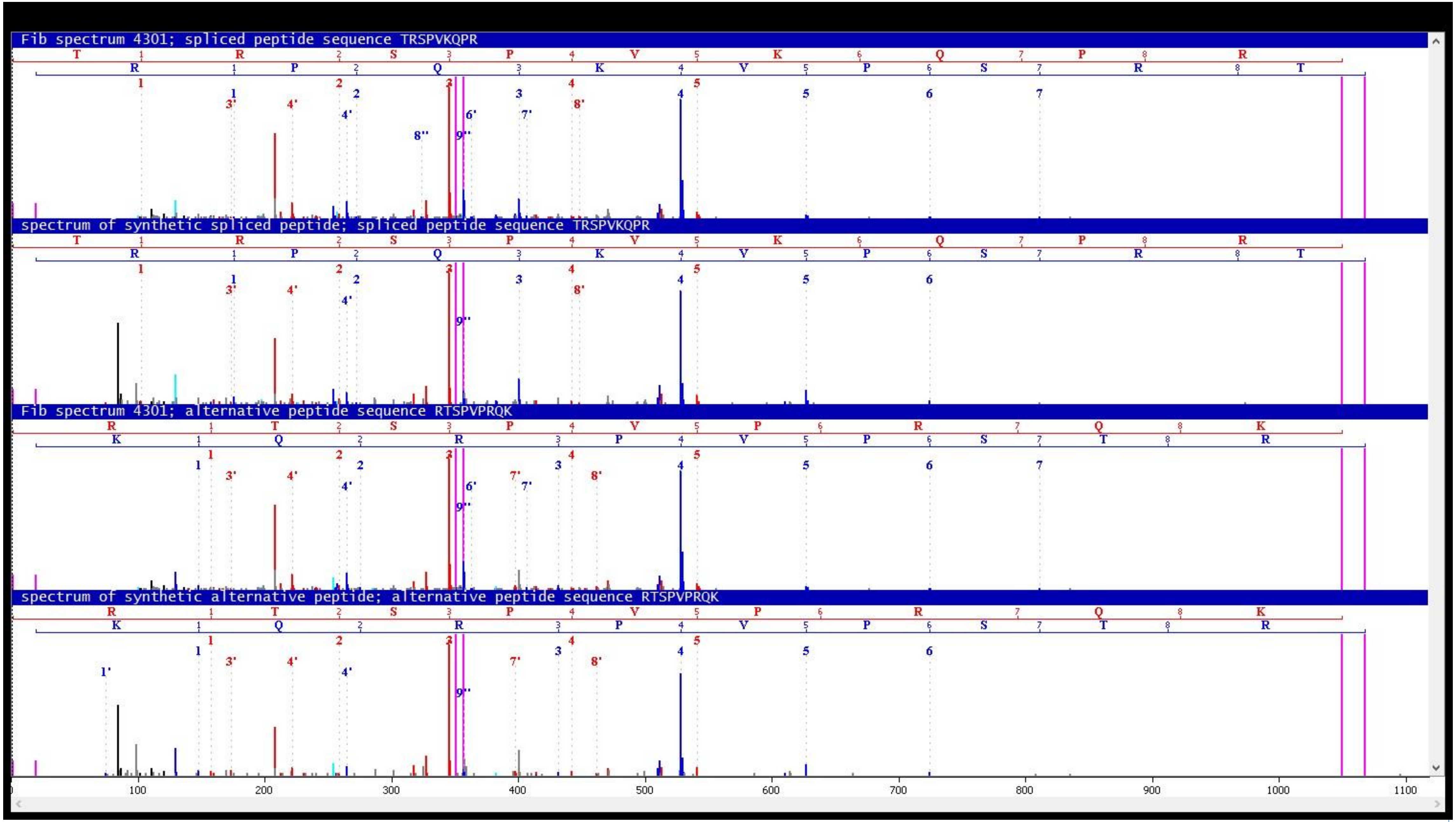
Fib spectrum 4301; spliced peptide sequence TRSPVKQPR
spectrum of synthetic spliced peptide; spliced peptide sequence TRSPVKQPR
Fib spectrum 4301; alternative peptide sequence RTSPVPRQK
spectrum of synthetic alternative peptide; alternative peptide sequence RTSPVPRQK
100
200
300
400
500
600
700
800
900
1000
1100

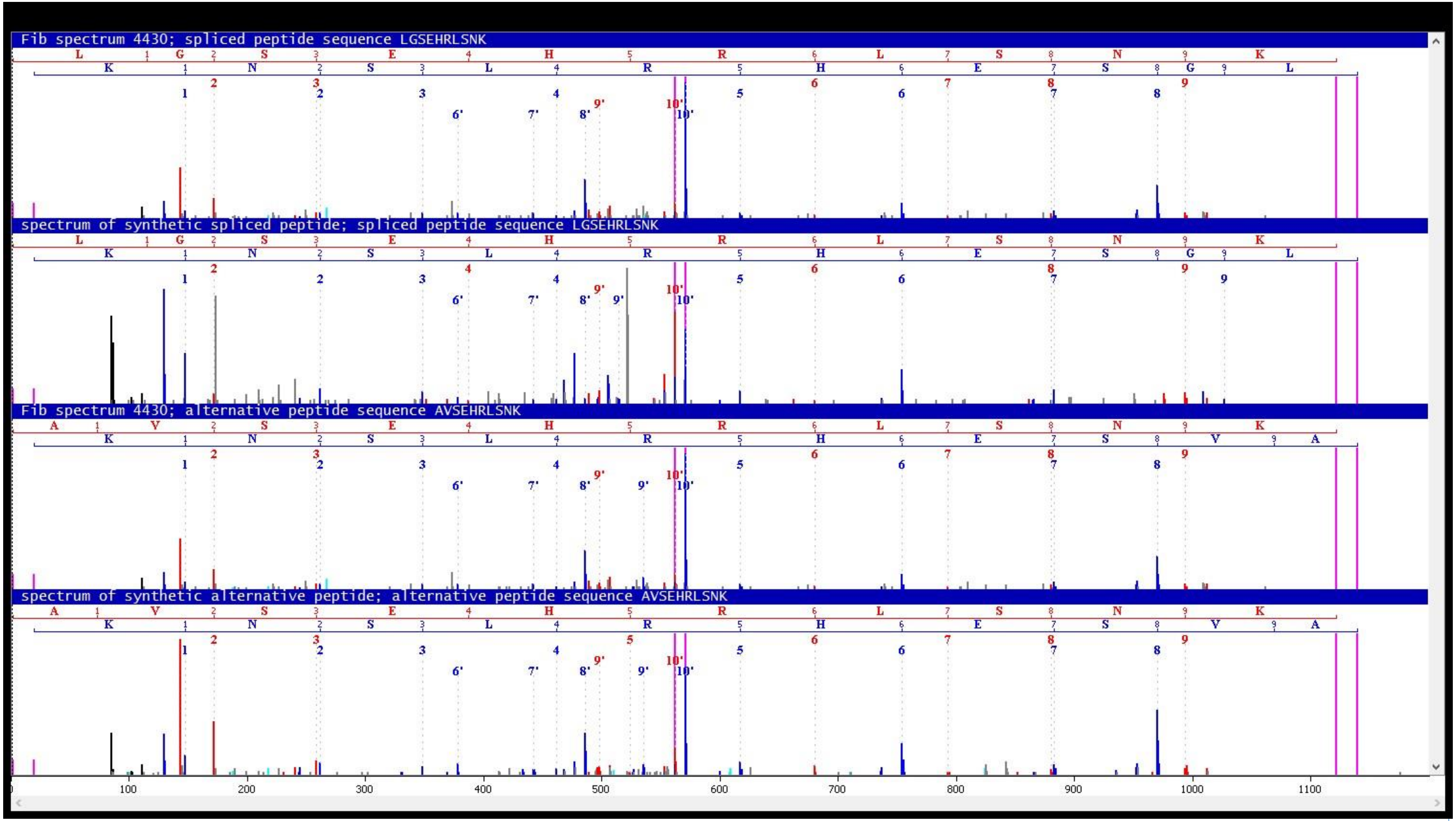

Fib spectrum 4430; spliced peptide sequence LGSEHRLSNK
spectrum of synthetic spliced peptide; spliced peptide sequence LGSEHRLSNK
Fib spectrum 4430; alternative peptide sequence AVSEHRLSNK
spectrum of synthetic alternative peptide; alternative peptide sequence AVSEHRLSNK
100
200
300
400
500
600
700
800
900
1000
1100

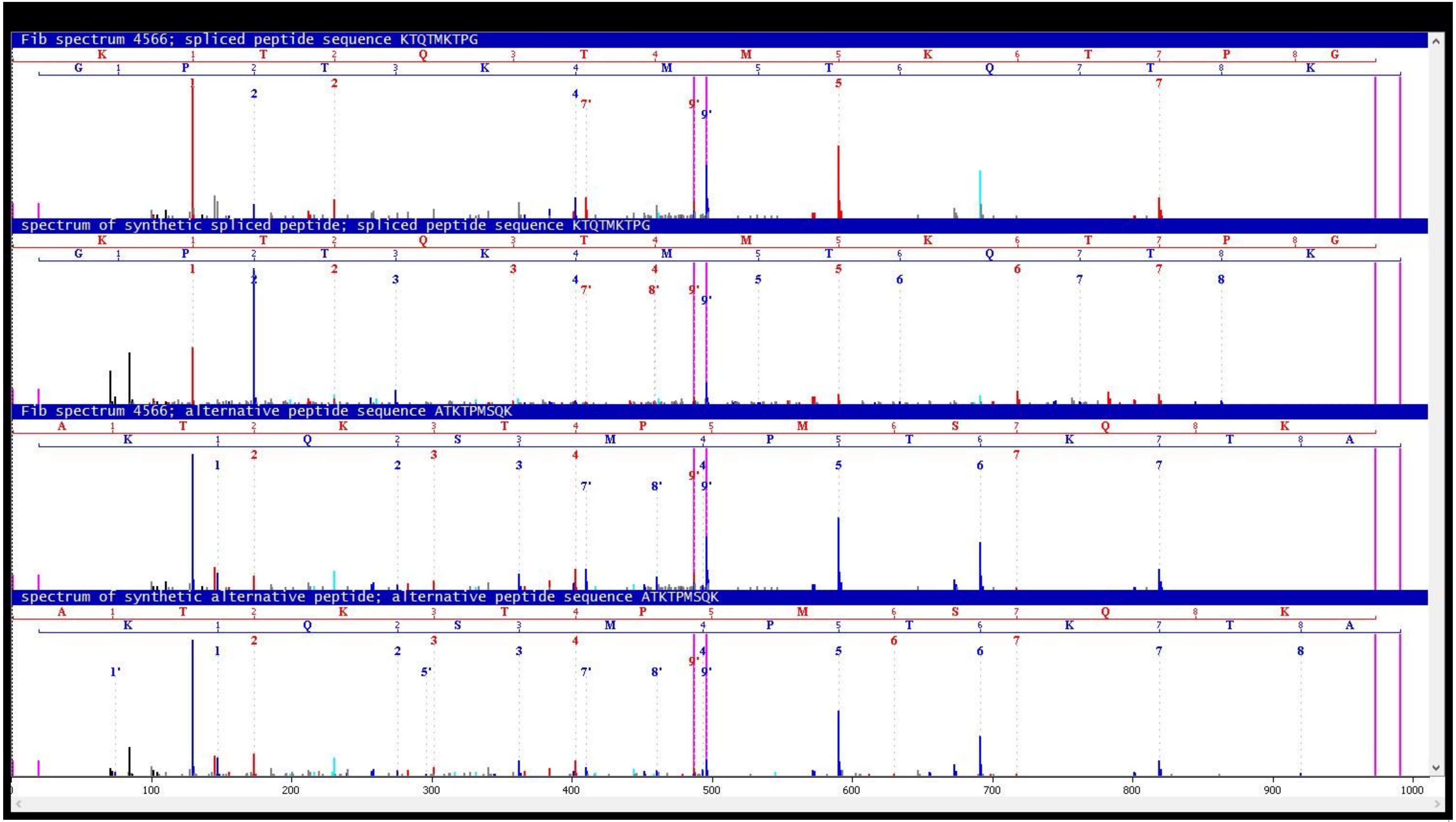
Fib spectrum 4566; spliced peptide sequence KTQTMKTPG
spectrum of synthetic spliced peptide; spliced peptide sequence KTQTMKTPG
Fib spectrum 4566; alternative peptide sequence ATKTPMSQK
spectrum of synthetic alternative peptide; alternative peptide sequence ATKTPMSQK
100
200
300
400
500
600
700
800
900
1000

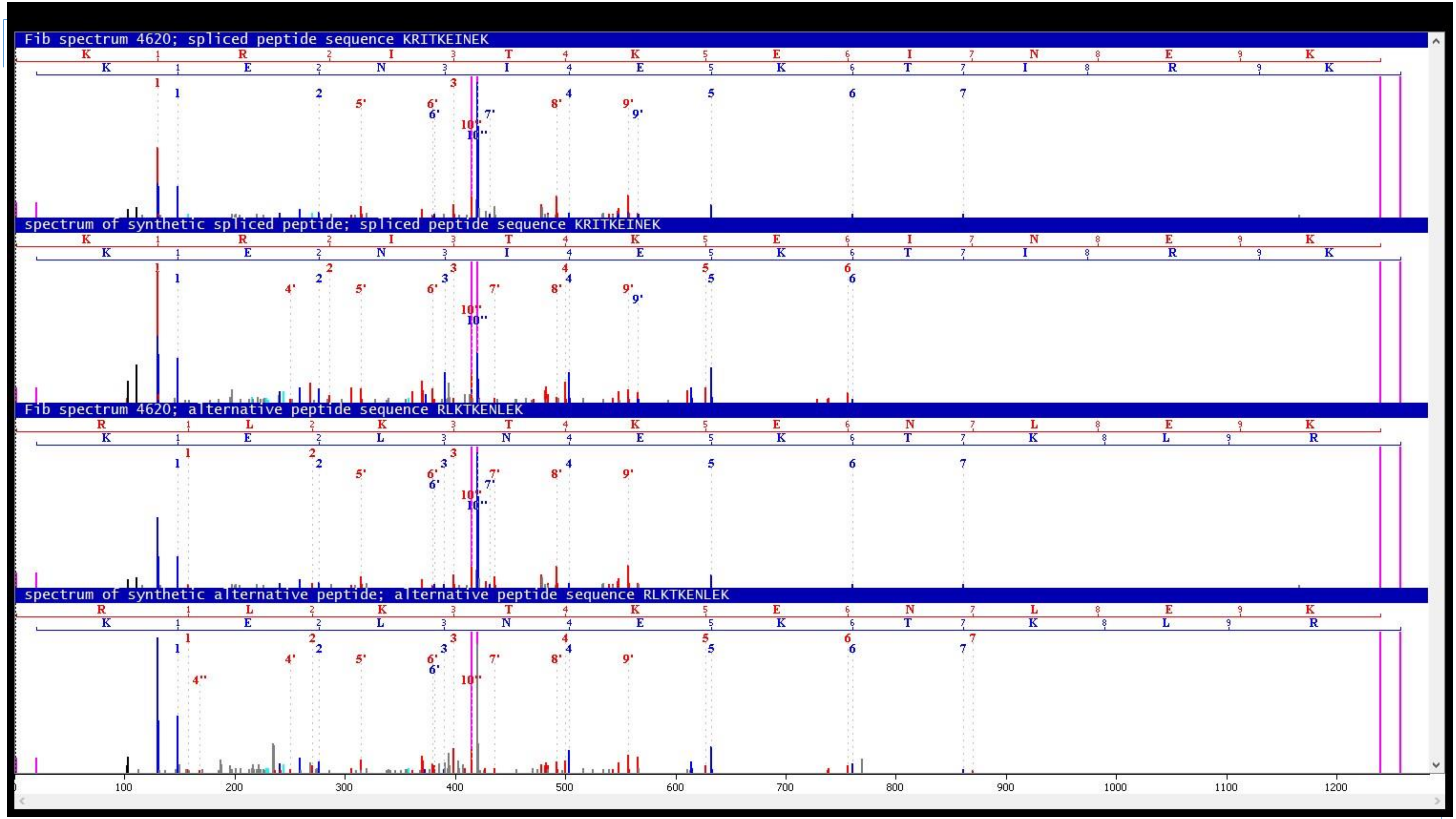
Fib spectrum 4620; spliced peptide sequence KRITKEINEK
spectrum of synthetic spliced peptide; spliced peptide sequence KRITKEINEK
Fib spectrum 4620; alternative peptide sequence RLKTKENLEK
spectrum of synthetic alternative peptide; alternative peptide sequence RLKTKENLEK
100
200
300
400
500
600
700
800
900
1000
1100
1200

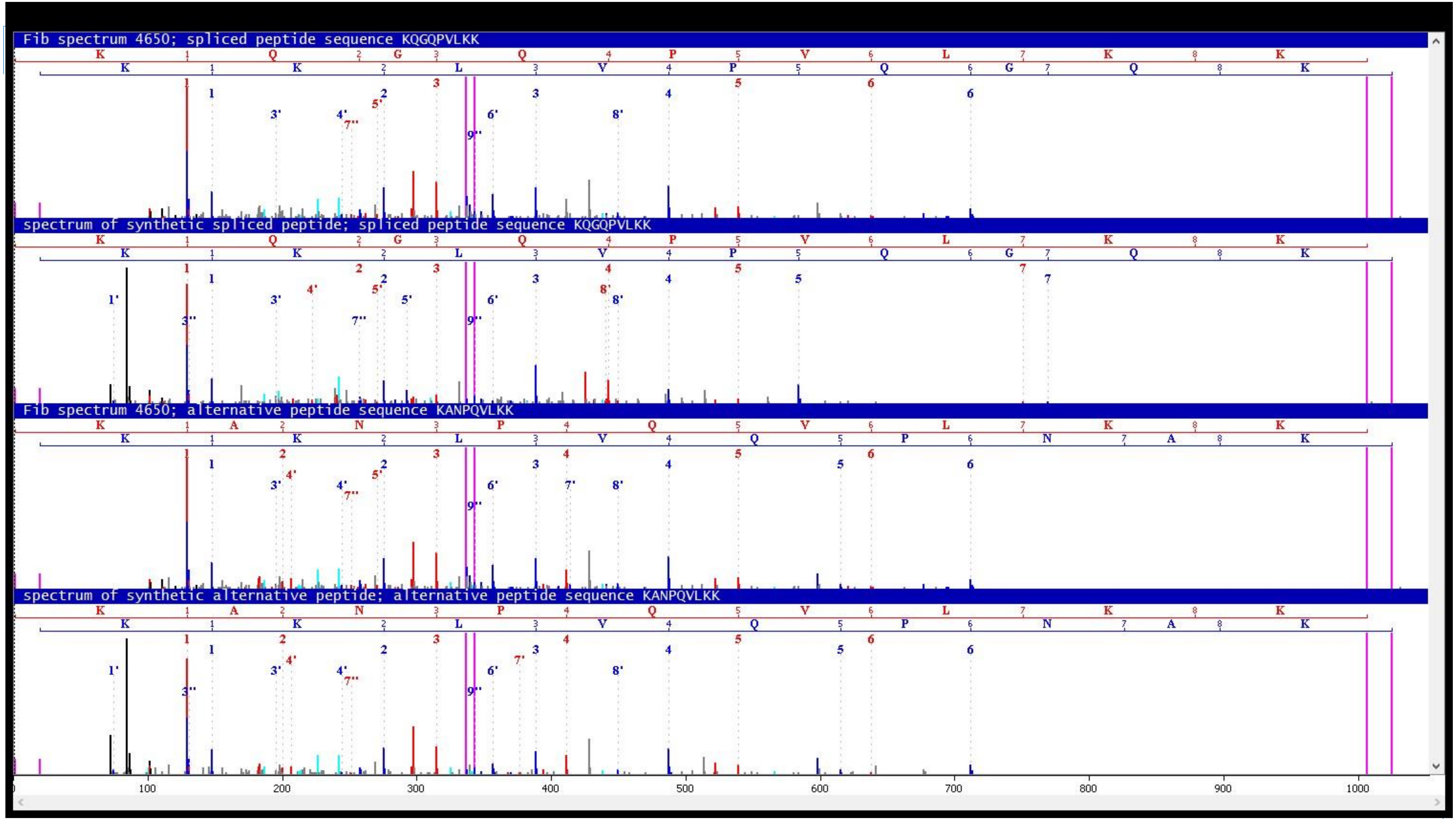

Fib spectrum 4650; spliced peptide sequence KQGQPVLKK
spectrum of synthetic spliced peptide; spliced peptide sequence KQGQPVLKK
Fib spectrum 4650; alternative peptide sequence KANPQVLKK
spectrum of synthetic alternative peptide; alternative peptide sequence KANPQVLKK
100
200
300
400
500
600
700
800
900
1000

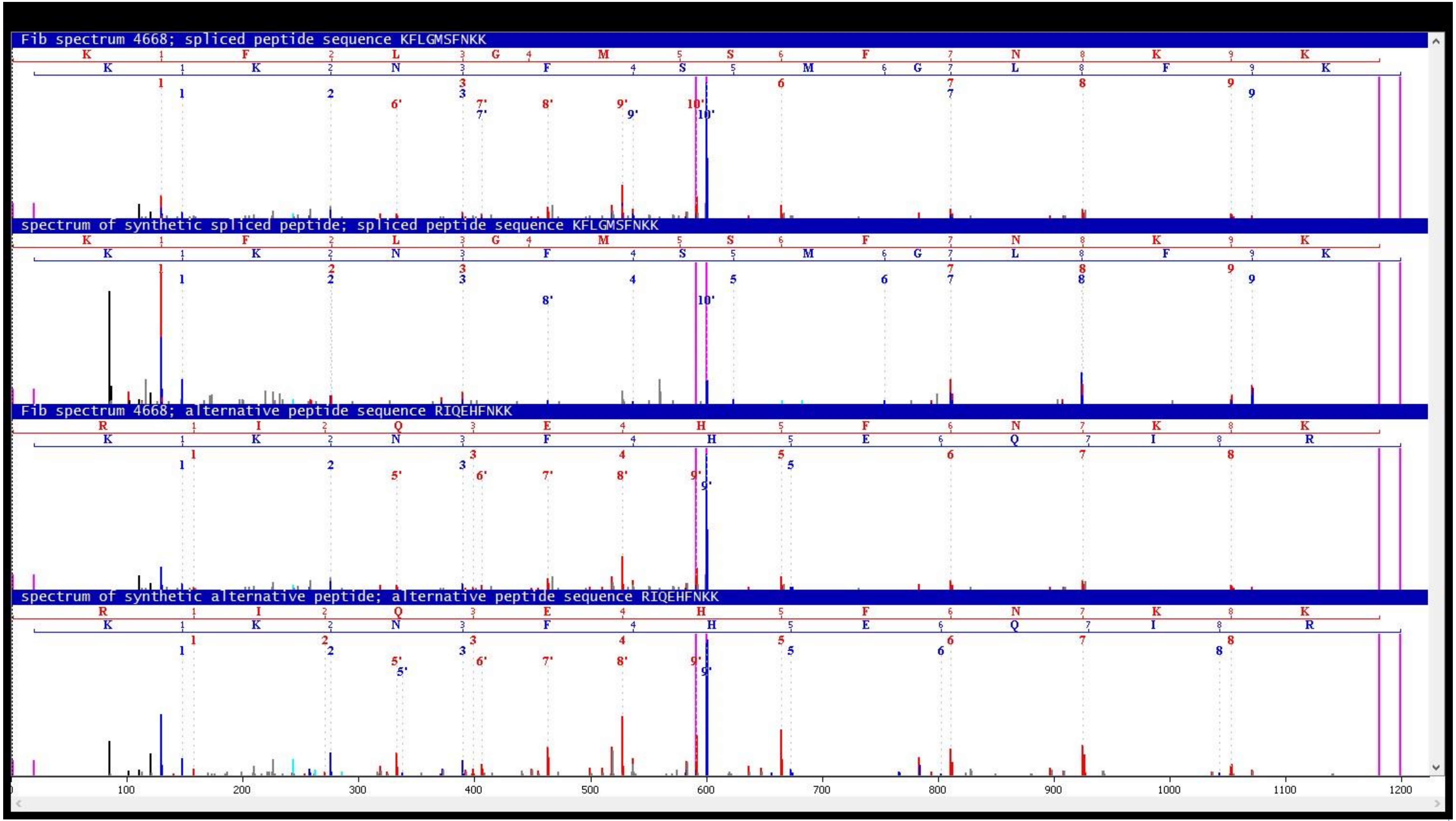
Fib spectrum 4668; spliced peptide sequence KFLGMSFNKK
spectrum of synthetic spliced peptide; spliced peptide sequence KFLGMSFNKK
Fib spectrum 4668; alternative peptide sequence RIQEHFNKK
spectrum of synthetic alternative peptide; alternative peptide sequence RIQEHFNKK
100
200
300
400
500
600
700
800
900
1000
1100
1200

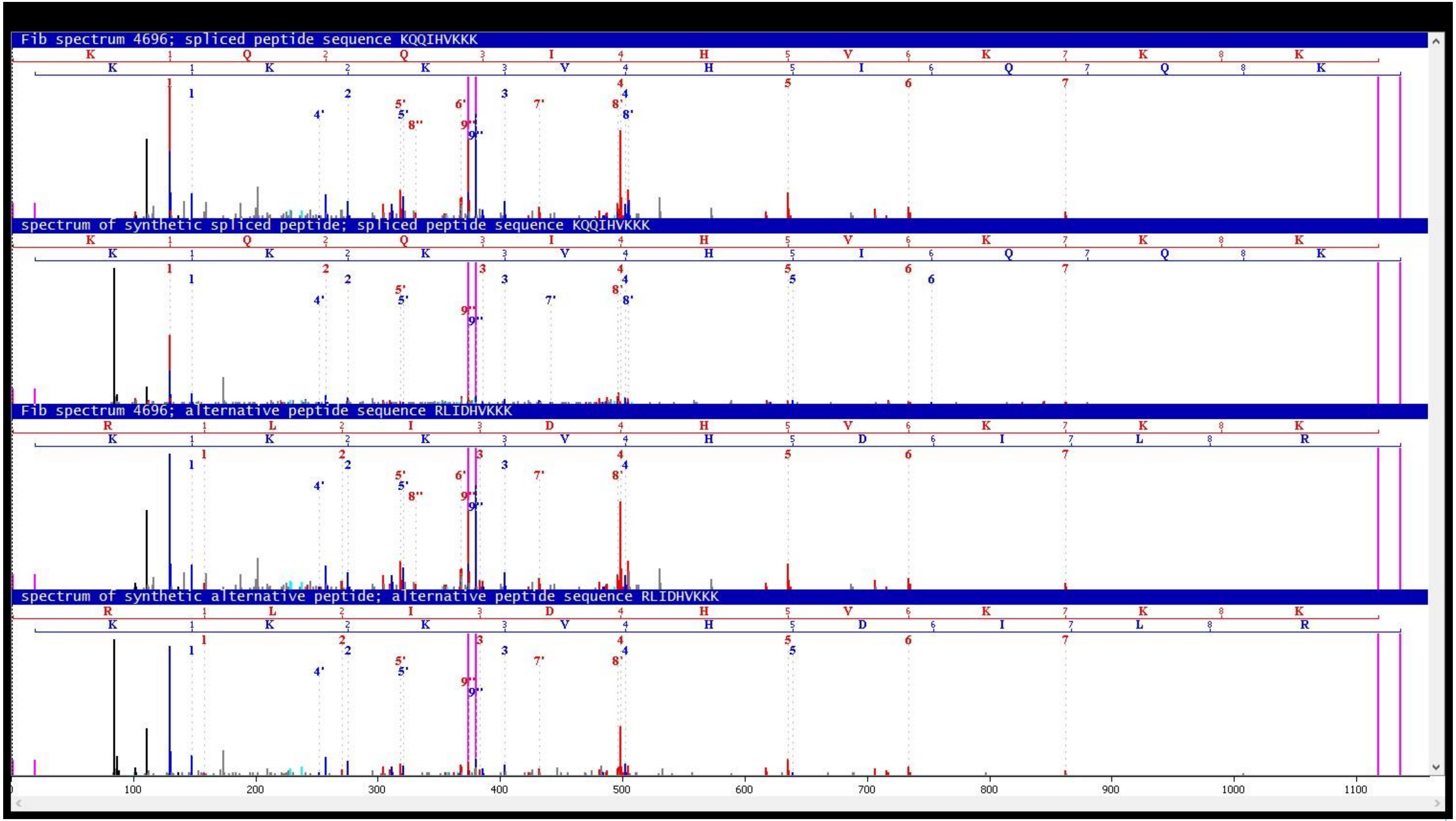

Fib spectrum 4696; spliced peptide sequence KQQIHVKKK
spectrum of synthetic spliced peptide; spliced peptide sequence KQQIHVKKK
Fib spectrum 4696; alternative peptide sequence RLIDHVKKK
spectrum of synthetic alternative peptide; alternative peptide sequence RLIDHVKKK
100
200
300
400
500
600
700
800
900
1000
1100

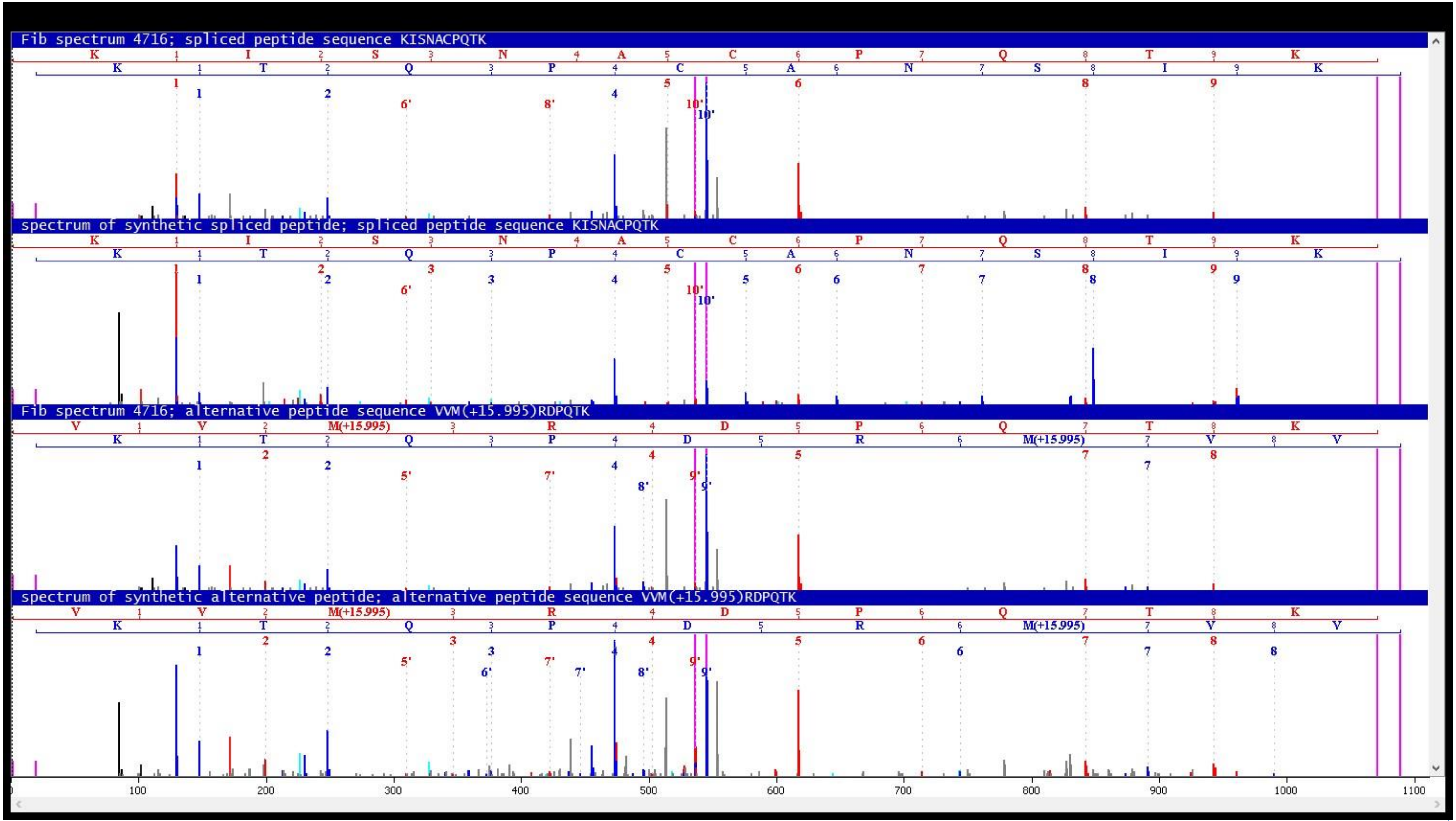

Fib spectrum 4716; spliced peptide sequence KISNACPQTK
spectrum of synthetic spliced peptide; spliced peptide sequence KISNACPQTK
Fib spectrum 4716; alternative peptide sequence VVM(+15.995)RDPQTK
spectrum of synthetic alternative peptide; alternative peptide sequence VVM(+15.995)RDPQTK
100
200
300
400
500
600
700
800
900
1000
1100

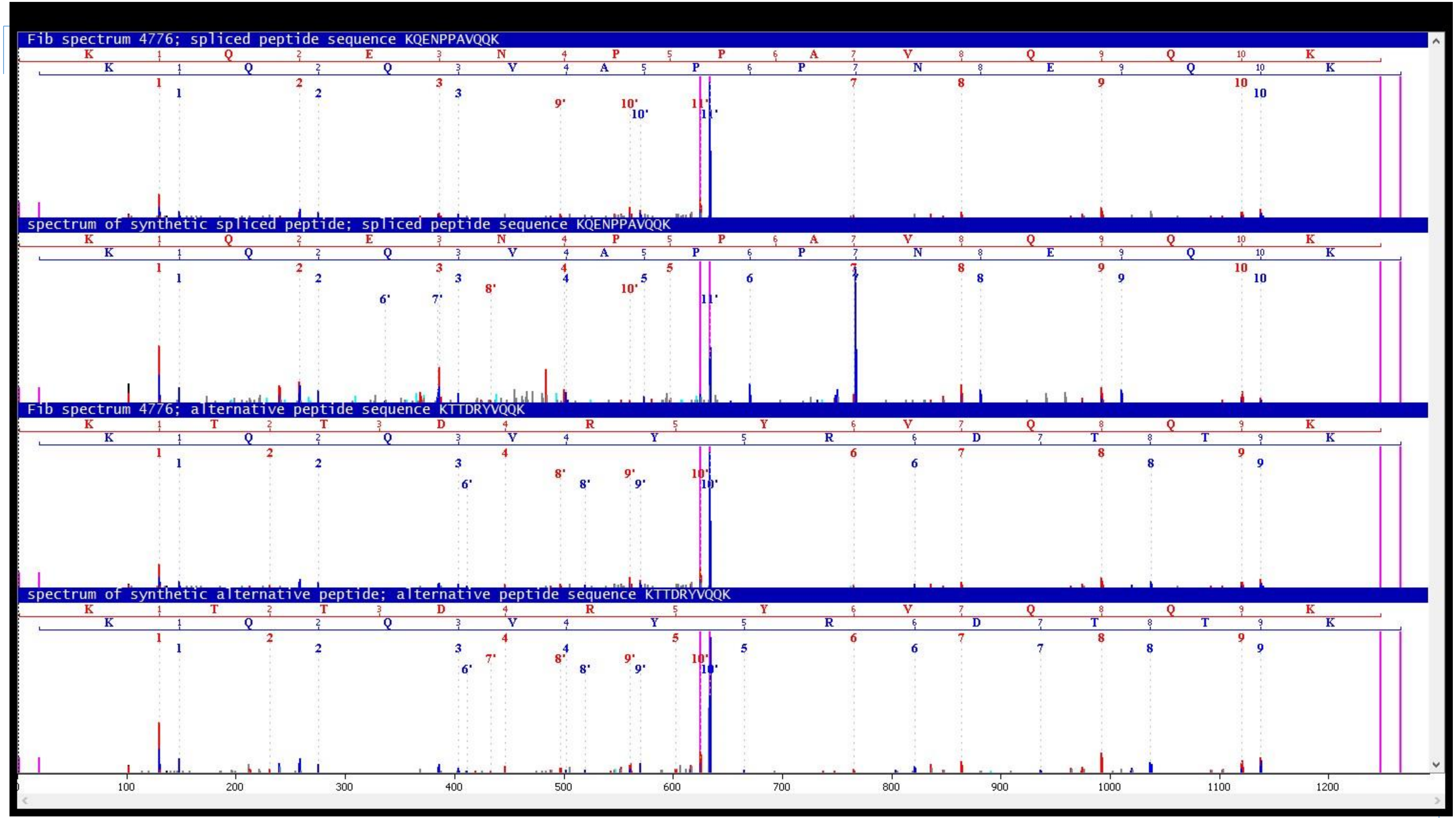
Fib spectrum 4776; spliced peptide sequence KQENPPAVQQK
spectrum of synthetic spliced peptide; spliced peptide sequence KQENPPAVQQK
Fib spectrum 4776; alternative peptide sequence KTTDRYVQQK
spectrum of synthetic alternative peptide; alternative peptide sequence KTTDRYVQQK
100
200
300
400
500
600
700
800
900
1000
1100
1200

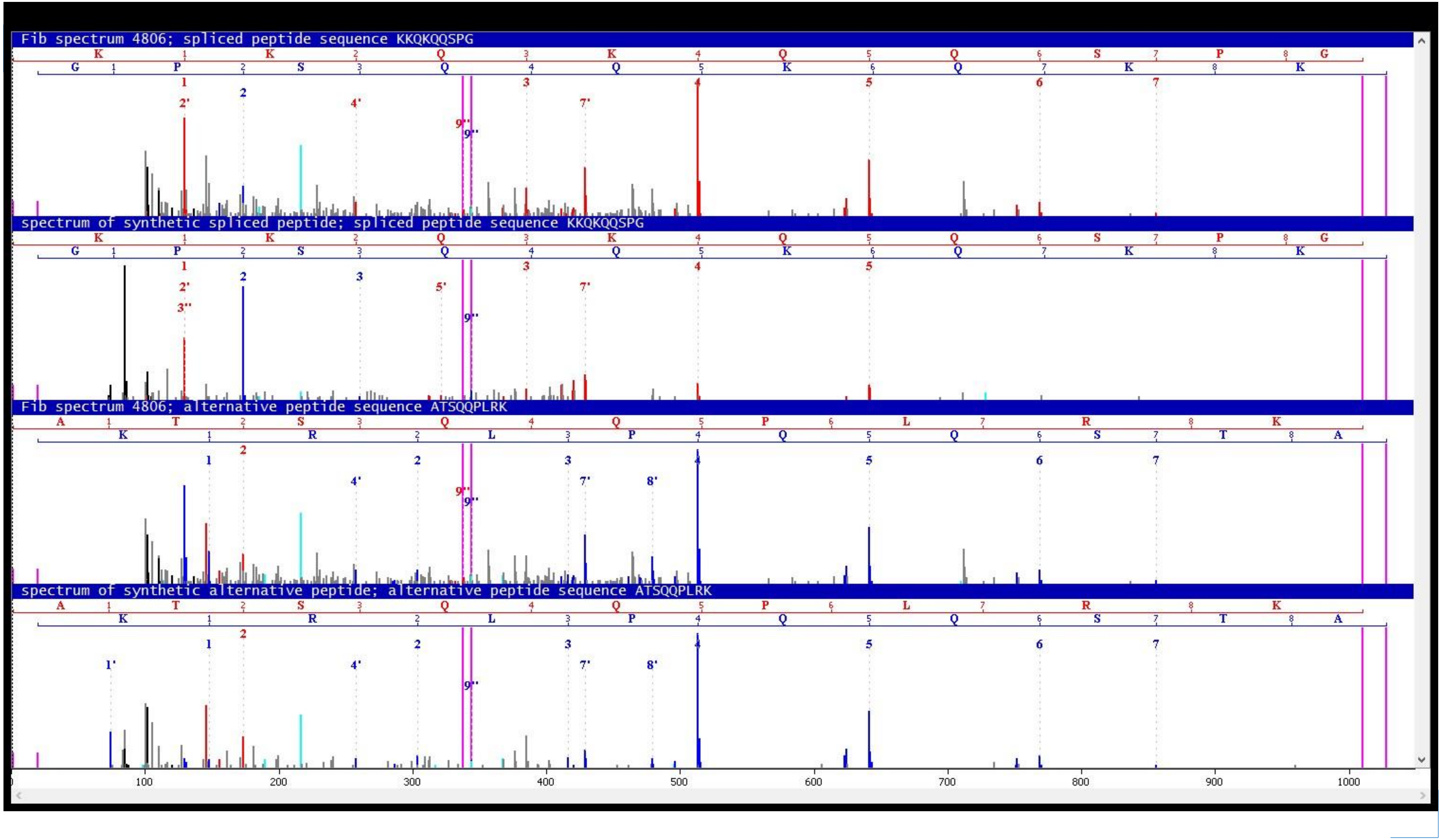
Fib spectrum 4806; spliced peptide sequence KKQKQQSPG
spectrum of synthetic spliced peptide; spliced peptide sequence KKQKQQSPG
Fib spectrum 4806; alternative peptide sequence ATSQQPLRK
spectrum of synthetic alternative peptide; alternative peptide sequence ATSQQPLRK
100
200
300
400
500
600
700
800
900
1000

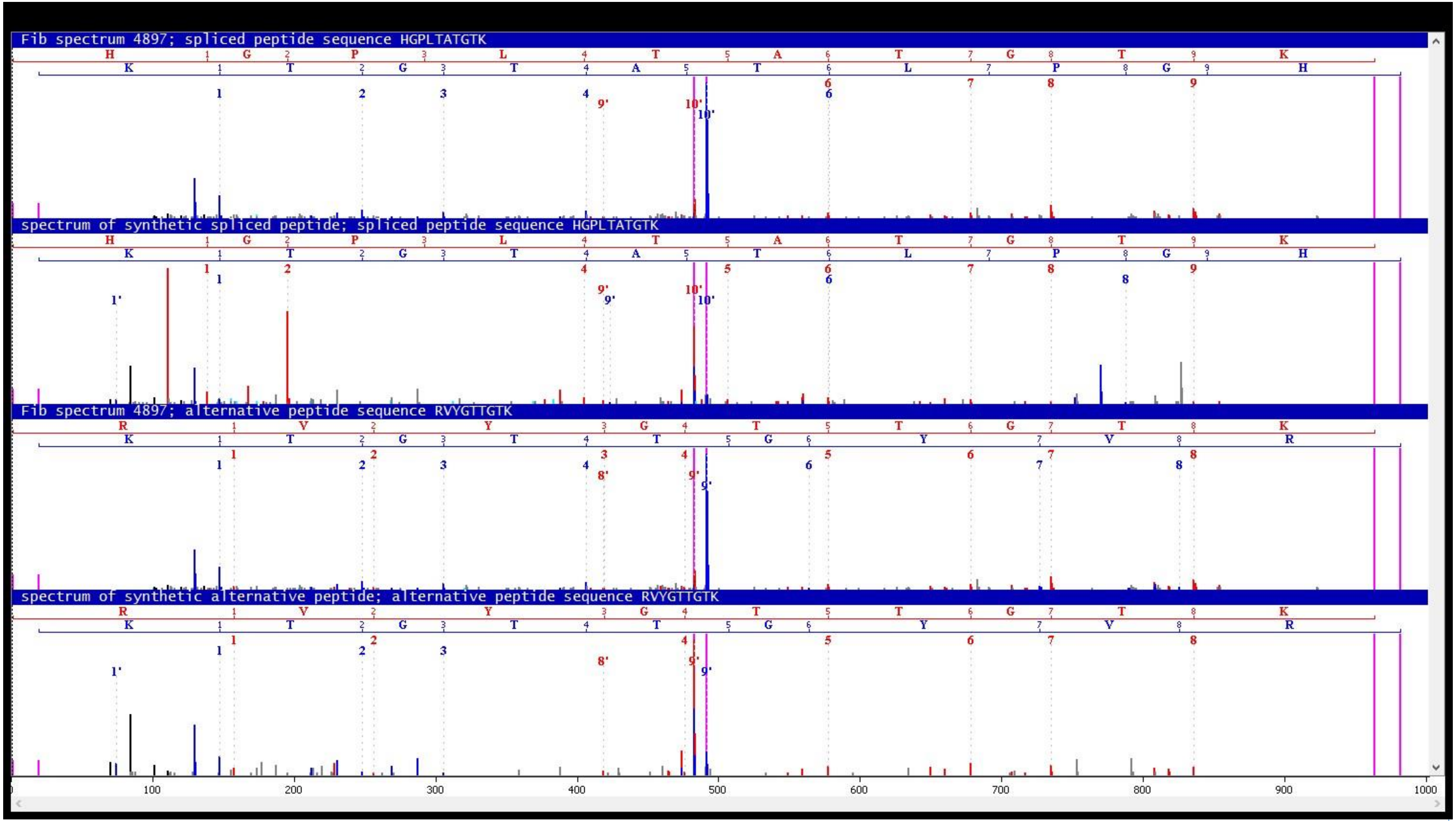
Fib spectrum 4897; spliced peptide sequence HGPLTATGTK
spectrum of synthetic spliced peptide; spliced peptide sequence HGPLTATGTK
Fib spectrum 4897; alternative peptide sequence RVYGTTGTK
spectrum of synthetic alternative peptide; alternative peptide sequence RVYGTTGTK
100
200
300
400
500
600
700
800
900
1000

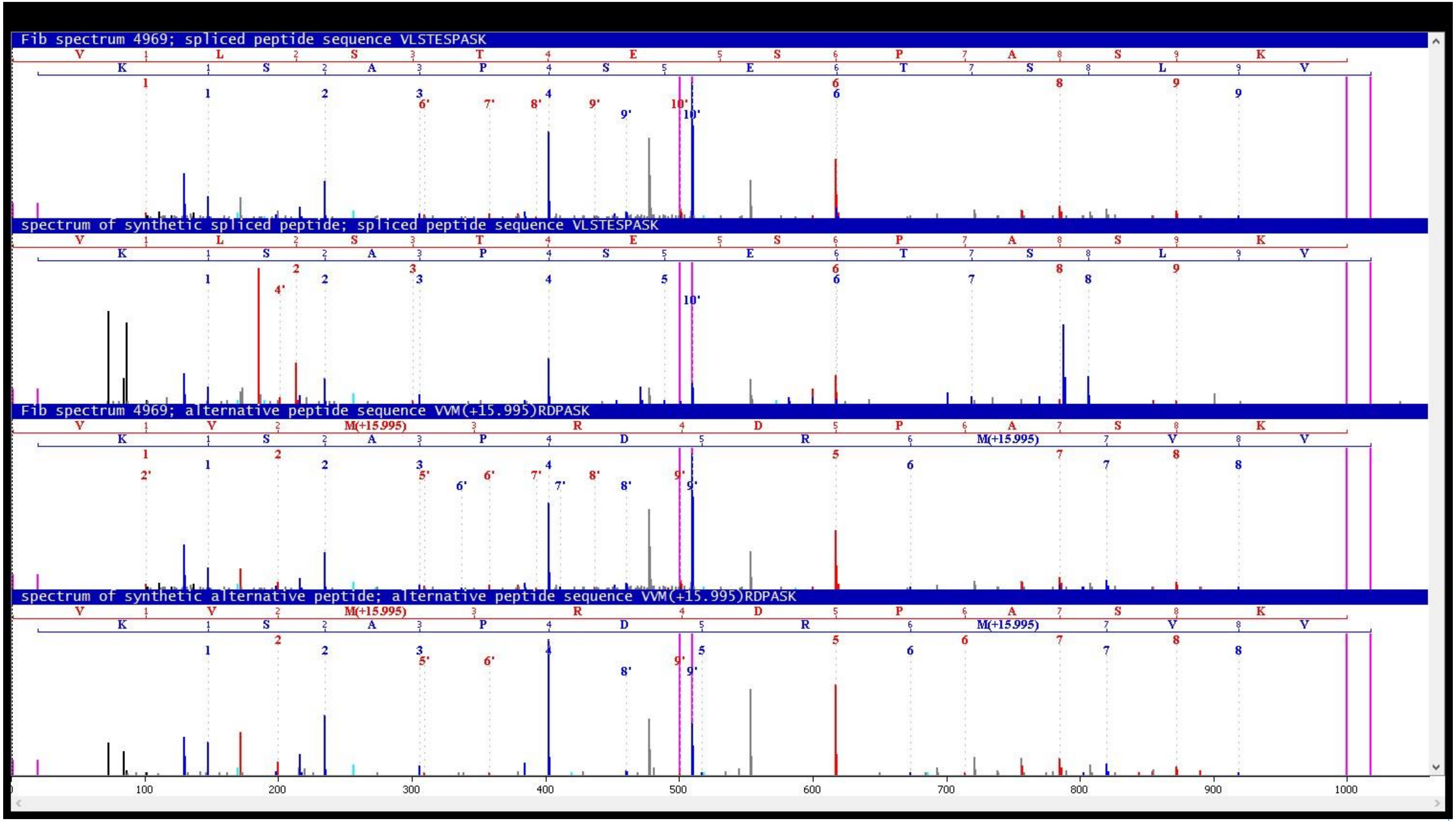
Fib spectrum 4969; spliced peptide sequence VLSTESPASK
spectrum of synthetic spliced peptide; spliced peptide sequence VLSTESPASK
Fib spectrum 4969; alternative peptide sequence VVM(+15.995)RDPASK
spectrum of synthetic alternative peptide; alternative peptide sequence VVM(+15.995)RDPASK
100
200
300
400
500
600
700
800
900
1000

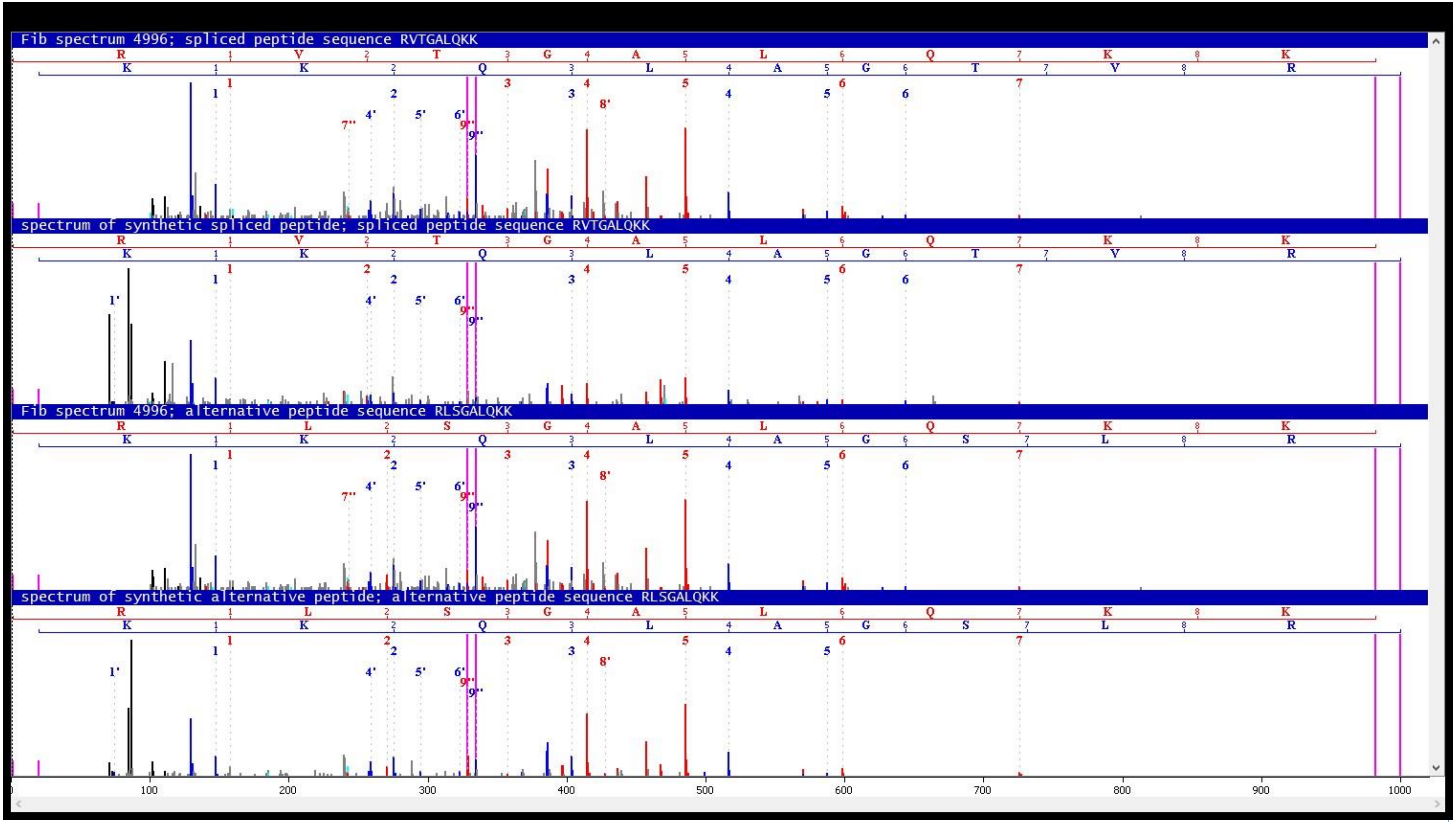
Fib spectrum 4996; spliced peptide sequence RVTGALQKK
spectrum of synthetic spliced peptide; spliced peptide sequence RVTGALQKK
Fib spectrum 4996; alternative peptide sequence RLSGALQKK
spectrum of synthetic alternative peptide; alternative peptide sequence RLSGALQKK
100
200
300
400
500
600
700
800
900
1000

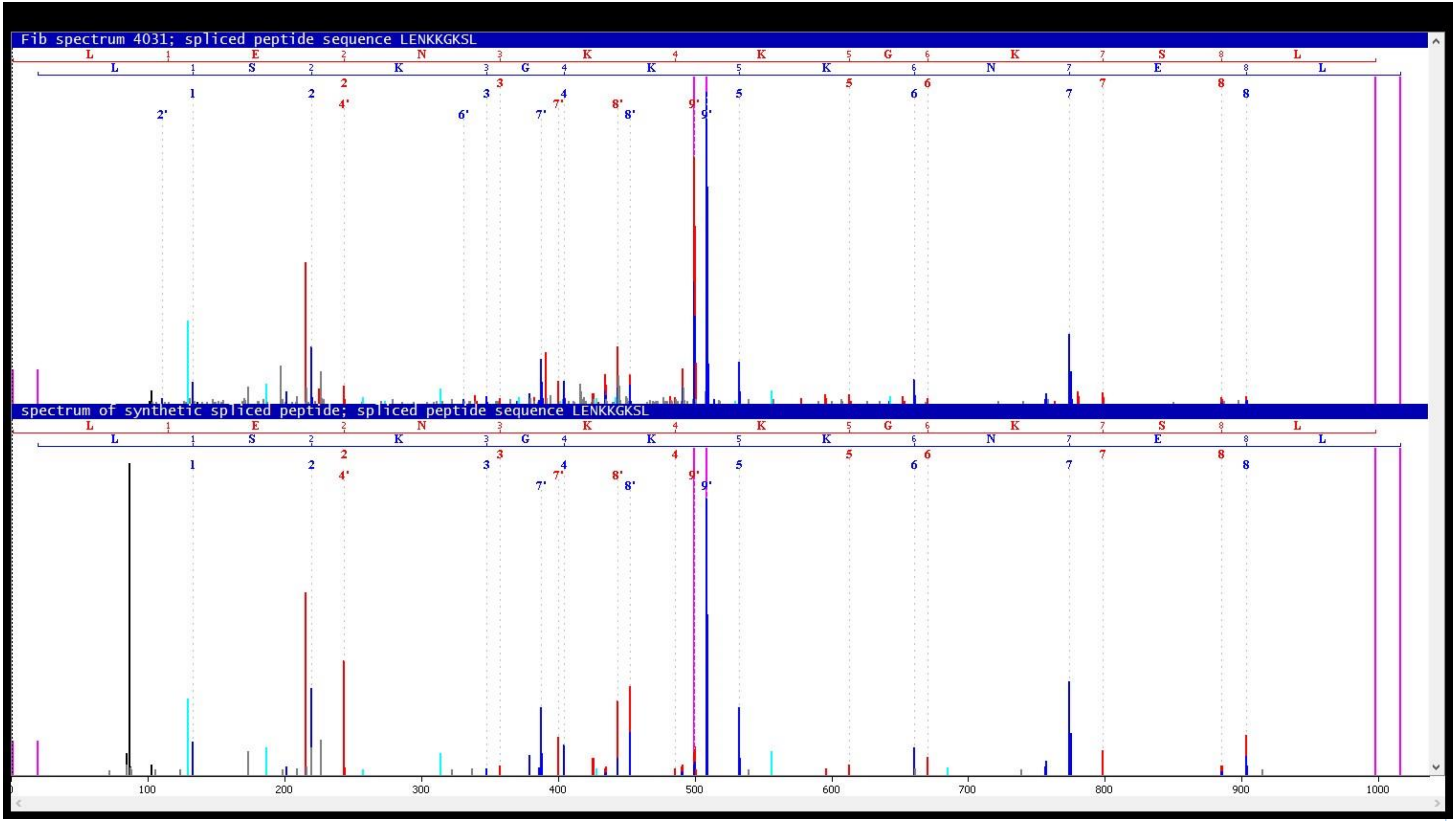
Fib spectrum 4031; spliced peptide sequence LENKKGKSL
spectrum of synthetic spliced peptide; spliced peptide sequence LENKKGKSL
100
200
300
400
500
600
700
800
900
1000

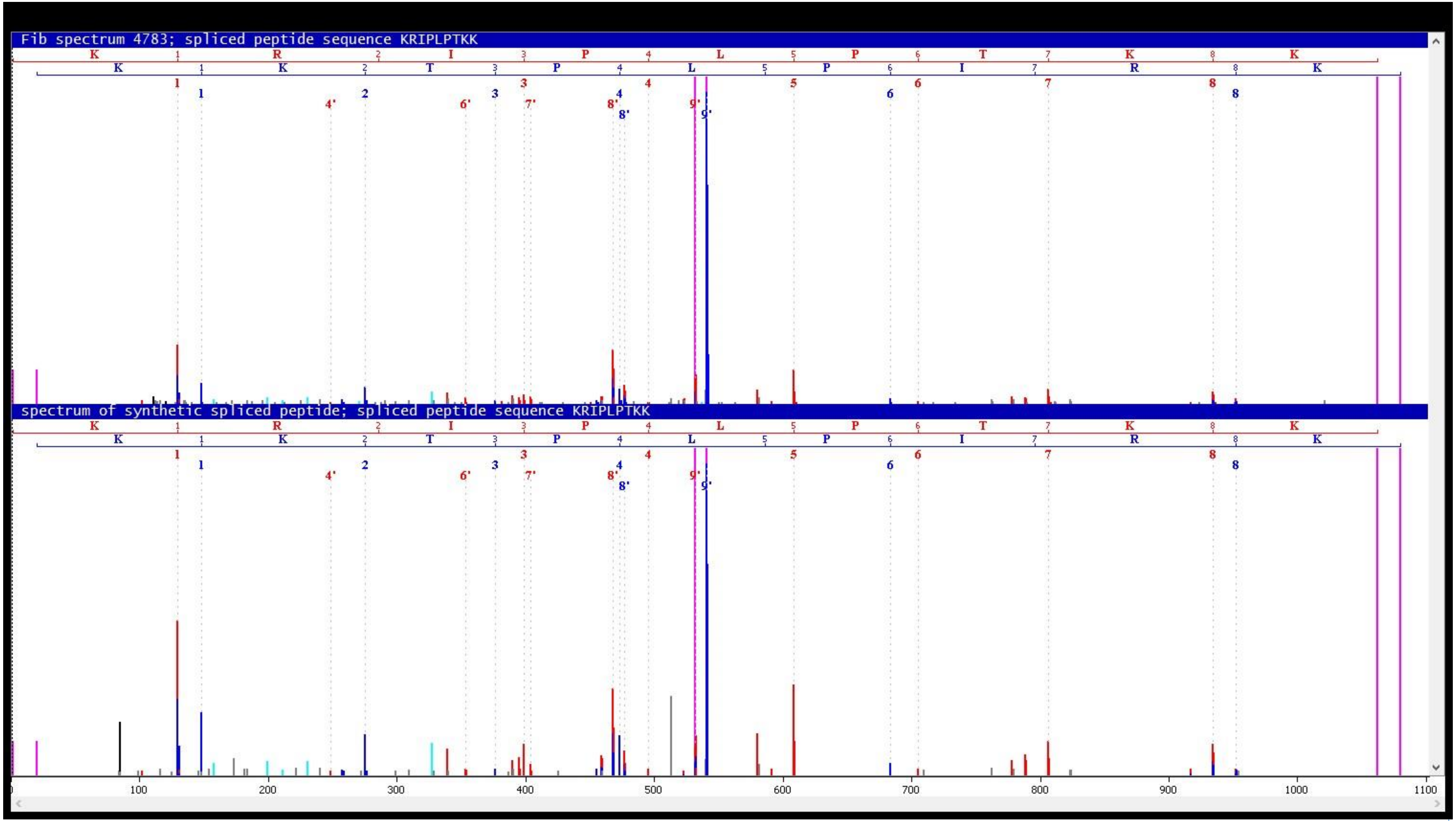
Fib spectrum 4783; spliced peptide sequence KRIPLPTKK
spectrum of synthetic spliced peptide; spliced peptide sequence KRIPLPTKK
100
200
300
400
500
600
700
800
900
1000
1100

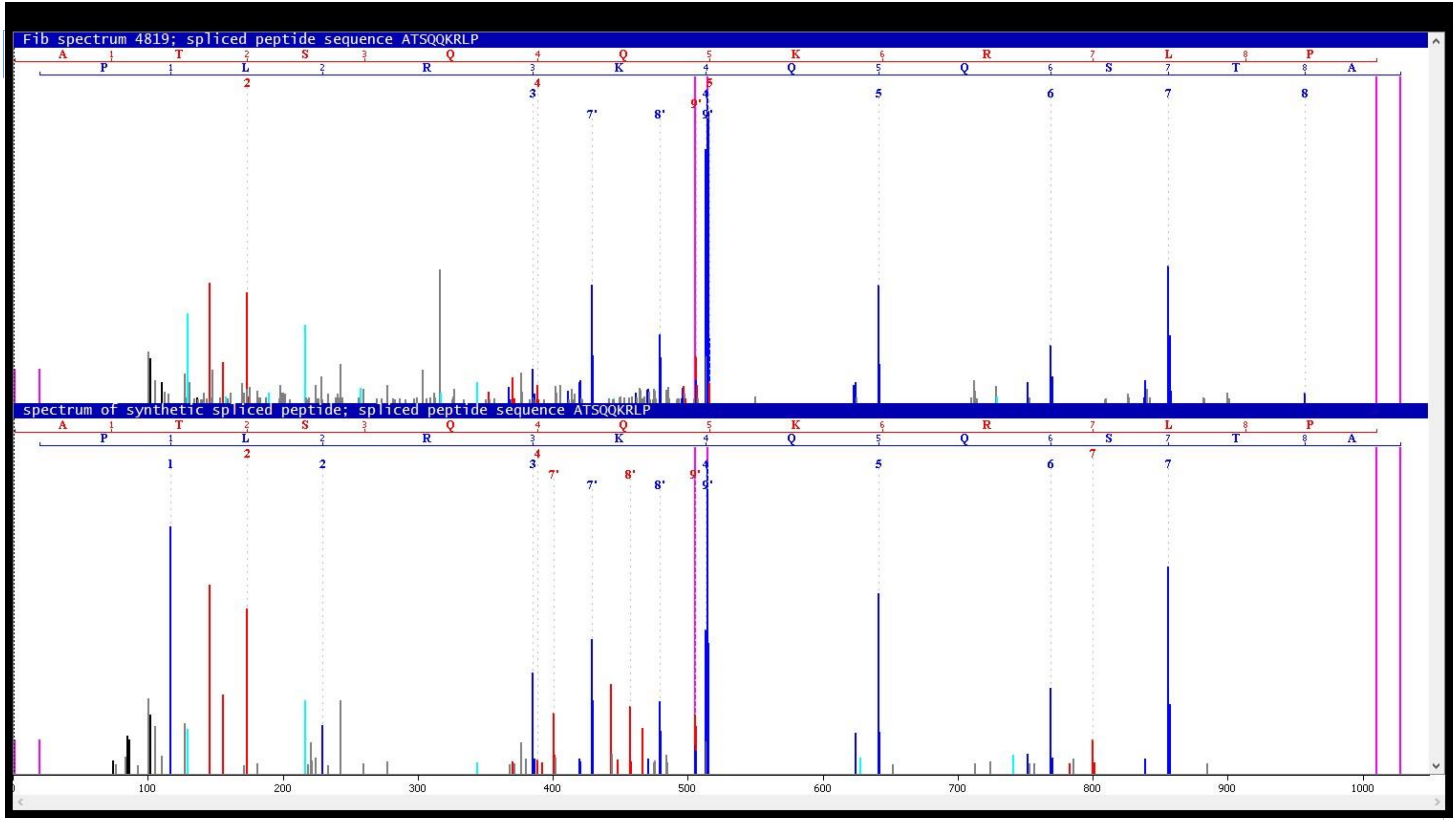

Fib spectrum 4819; spliced peptide sequence ATSQQKRLP
spectrum of synthetic spliced peptide; spliced peptide sequence ATSQQKRLP
100
200
300
400
500
600
700
800
900
1000

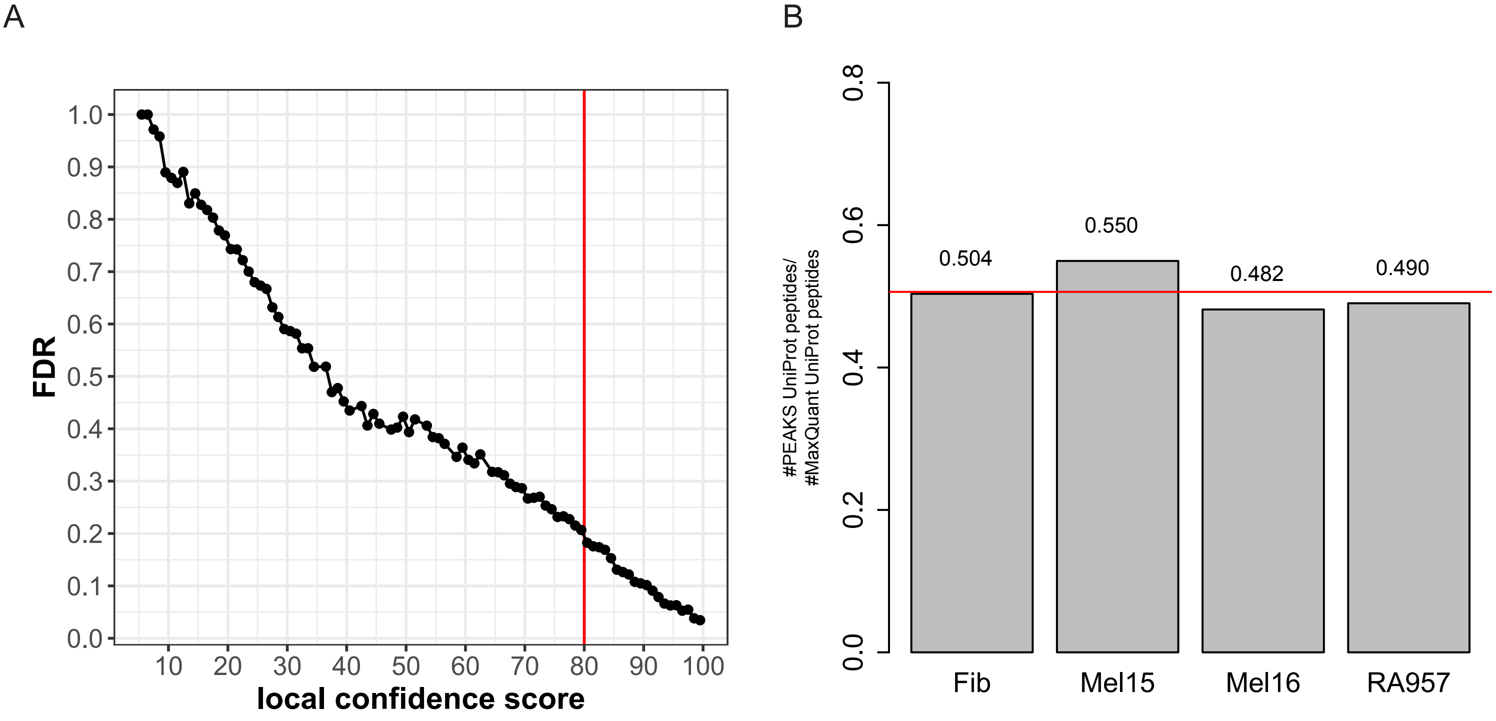


Supplementary Figure 5: A) Estimation of the error rate in the PEAKS results. This figure shows the fraction of Mel15 spectra, where the peptide sequence identified by PEAKS differed from the sequence identified by MaxQuant for the UniProt peptides, as a function of the PEAKS local confidence score. Since the Max-Quant FDR is in agreement with the Comet FDR and Mascot FDR for the UniProt peptides (Figure 1C), we assume that the error by MaxQuant is indeed about 1% and most of the sequence differences are caused by erroneous PEAKS matches. Therefore, the rate of differing sequences can be used as a rough proxy for the PEAKS FDR. PEAKS shows a FDR of about 20% at a local confidence score of 80 (red line). B) Rate of UniProt HLA peptides identified by PEAKS at a local confidence score of 80 compared to MaxQuant UniProt peptides at spectrum level of FDR 1% for the different samples.

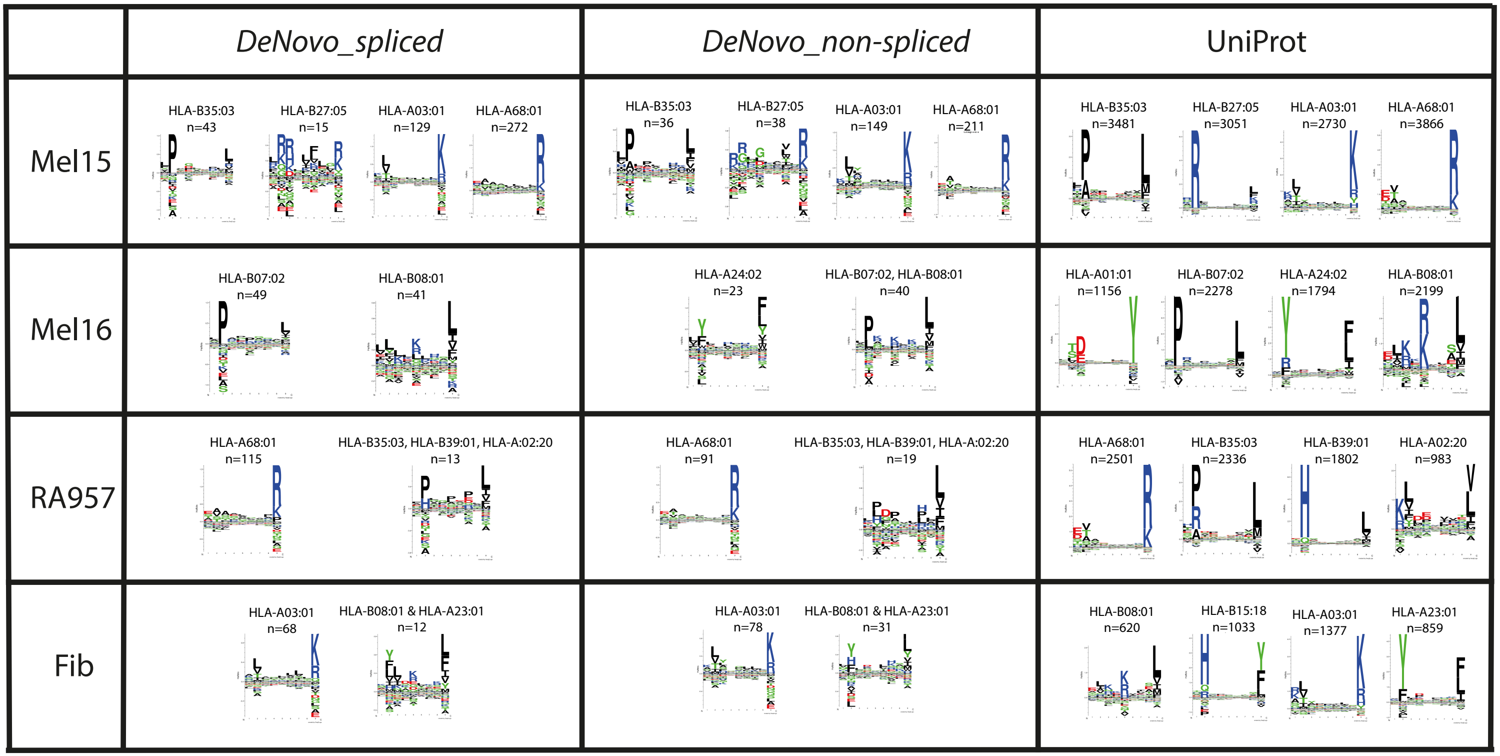


Supplamentary Figure 6: Sequence motifs for the 9-mer *DeNovo_spliced* peptides, *DeNovo_non-spliced* peptides and UniProt peptides identified by MaxQuant in Mel15, Mel16, RA957 and Fib samples.

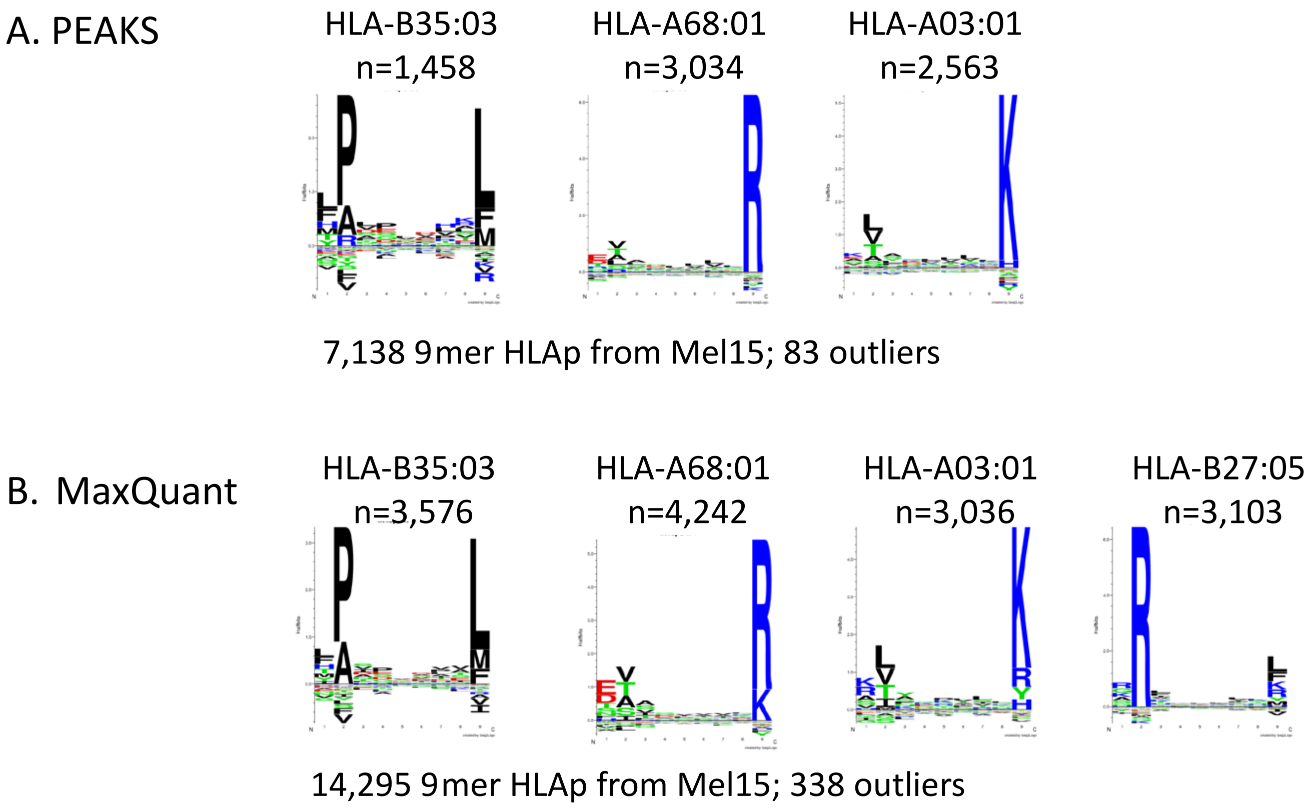


Supplementary Figure 7: A) Gibbs clustering analysis of Mel15 HLA-Ip identified as UniProt hits by PEAKS reveals three dominant clusters referring to HLA-B35:03, HLA-A68:01 and HLA-A03:01. B) Gibbs clustering analysis of HLA-Ip identified as UniProt hits by MaxQuant from the same MS data reveals four dominant clusters, including the motif of HLA-B27:05.
